# Global remodeling of ADP-ribosylation by PARP1 suppresses influenza A virus infection

**DOI:** 10.1101/2024.09.19.613696

**Authors:** Zhenyu Zhang, Isabel Uribe, Kaitlin A. Davis, Robert Lyle McPherson, Gloria P Larson, Mohsen Badiee, Vy Tran, Mitchell P. Ledwith, Elizabeth Feltman, Shuǐqìng Yú, Yíngyún Caì, Che-Yuan Chang, Xingyi Yang, Zhuo Ma, Paul Chang, Jens H Kuhn, Anthony K. L. Leung, Andrew Mehle

**Author notes:** These authors contributed equally (Zhenyu Zhang, Isabel Uribe).

## Abstract

ADP-ribosylation is a highly dynamic and fully reversible post-translational modification performed by poly(ADP-ribose) polymerases (PARPs) that modulates protein function, abundance, localization and turnover. Here we show that influenza A virus infection causes a rapid and dramatic upregulation of global ADP-ribosylation that inhibits viral replication. Mass spectrometry defined for the first time the global ADP-ribosylome during infection, creating an infection-specific profile with almost 4,300 modification sites on ∼1,080 host proteins, as well as over 100 modification sites on viral proteins. Our data indicate that the global increase likely reflects a change in the form of ADP-ribosylation rather than modification of new targets. Functional assays demonstrated that modification of the viral replication machinery antagonizes its activity and further revealed that the anti-viral activity of PARPs and ADP-ribosylation is counteracted by the influenza A virus protein NS1, assigning a new activity to the primary viral antagonist of innate immunity. We identified PARP1 as the enzyme producing the majority of poly(ADP-ribose) present during infection. Influenza A virus replicated faster in cells lacking PARP1, linking PARP1 and ADP-ribosylation to the anti-viral phenotype. Together, these data establish ADP-ribosylation as an anti-viral innate immune-like response to viral infection antagonized by a previously unknown activity of NS1.

## INTRODUCTION

Diverse cellular stresses, including infection, reprogram the post-translational modifications of proteins, rapidly altering existing cellular landscapes to promote survival of the host. These modifications dynamically modulate target protein function, abundance, turnover, and localization, and negatively or positively regulate host responses and immune activation during infection^1–4^. Multiple lines of evidence have recently implicated ADP-ribosylation, along with ADP-ribosyltransferases (commonly known as poly(ADP-ribose) polymerases [PARPs]^5^), as an important part of this anti-pathogen response^6–9^. Many PARP genes are induced by interferon (IFN) signaling, suggesting a role in the response to pathogens^10–14^. Moreover, multiple *PARP* genes show signs of strong positive selection or have undergone gene loss or duplication m—processes that are hallmarks of evolutionary conflicts at the virus:host interface^15,16^. Successful viruses therefore need to antagonize, evade, or co-opt PARP-mediated cellular countermeasures to ensure their fitness. Consequently, multiple positive-sense RNA viruses are known to encode enzymes that remove ADP-ribose (ADPr) from proteins^6,8^. PARPs and ADP-ribosylation are an under-appreciated arm of cellular anti-viral responses.

PARPs are a family of enzymes involved in post-translational modifications affecting diverse biological functions, including DNA repair, gene regulation, and modulation of immune responses^17^. Of the 17 members of the human PARP family, 15 are catalytically active and transfer ADPr moieties from nicotinamide adenine dinucleotide onto target proteins or nucleic acids^18,19^. The extent and form of ADP-ribosylation can exert differential effects on the modified protein^17,20,21^. PARPs can attach a single ADPr to a target protein [mono(ADP-ribosyl)ation or MARylation] or multimeric chains of ADPr [poly(ADP-ribosyl)ation or PARylation]. ADP-ribosylation is dynamic and reversible. PARPs add ADPr to targets while ADP-ribosylhydrolases remodel PAR chains or completely remove ADPr^22–24^.

Influenza is a recurring global threat to animals, including humans. Viral replication is entirely dependent upon cellular cofactors, with influenza viruses usurping host factors to support replication while avoiding or evading anti-viral factors. PARP1 and PARP13 have previously been implicated as cellular factors that regulate influenza A virus replication. PARP1 binds the viral polymerase and may aid its function^25–27^. Whether the effects of PARP1 are mediated via ADP-ribosylation remains unclear. PARP13 (ZC3HAV1, also called zinc-finger anti-viral protein [ZAP]) inhibits viral replication, even though it is catalytically inactive^18,28,29^.

Here we show that influenza A virus infection causes a dramatic upregulation of global ADP-ribosylation that inhibits viral replication. We define for the first time the global ADP-ribosylome during influenza A virus infection, mapping thousands of modifications on viral and host proteins at single amino acid resolution. Infection induces a significant increase in PARylation to already modified proteins. Modifications on the viral replication machinery disable its activity, whereas the anti-viral activity of ADP-ribosylation is suppressed by the influenza A virus protein NS1. PARP1 is identified as the primary enzyme responsible for infection-induced PARylation and its anti-viral activity. Thus, ADP-ribosylation tempers viral replication as a key component of cellular anti-viral responses to influenza A virus, a process that is counteracted by the viral NS1 protein.

## RESULTS

### PARPs and ADP-ribosylation antagonize influenza A virus infection and replication

To identify host factors that impact influenza A virus, we performed a gene correlation analysis linking expression of cellular genes to susceptibility to infection. We previously characterized pro-viral factors that enhance infection^30^, and hence focus here on high-confidence anti-viral candidates that suppress infection. Among the putative anti-viral factors with the strongest negative correlation scores were interferon-induced transmembrane proteins (IFITMs) 1, 2, and 3 (Figure 1A). IFITMs strongly inhibit replication of several RNA viruses^31^, providing confidence in the ability of our screen to detect anti-viral factors.

**Figure 1.**
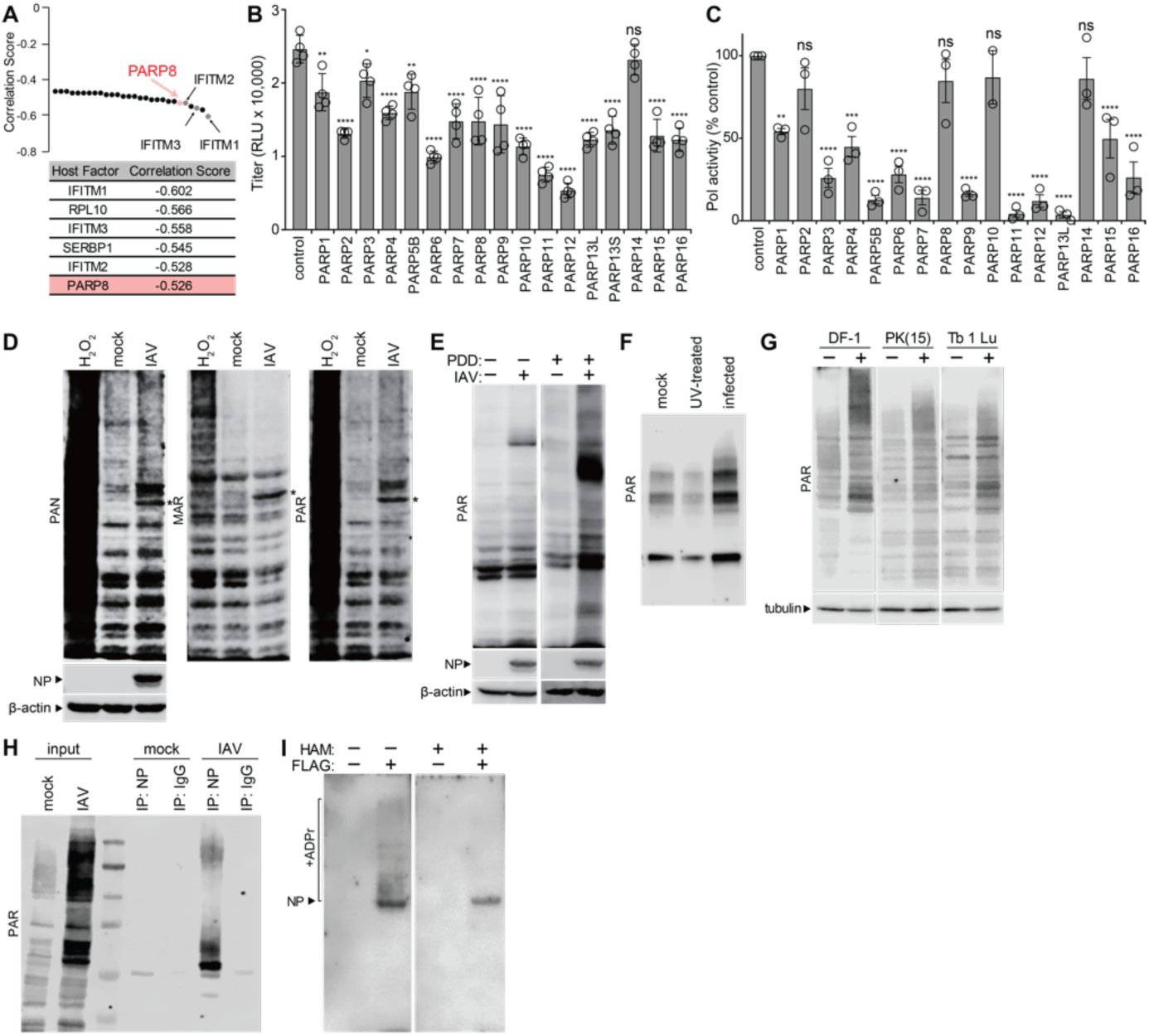
Anti-viral activity of PARPs and virus-induced ADP-ribosylation. **(A)** Gene correlation analysis identifies PARP8 as anti-viral factor. Top putative anti-viral hits from the gene correlation screen and their correlation scores, selected from our previously published full results^30^. **(B-C)** PARPs interfere with influenza A virus WSN. (B) Viral titers were measured after infection of cells expressing a member of the PARP family. (C) Polymerase activity assays were performed in the presence or absence of the indicated PARP expression. **(D)** Human lung A549 cells were infected with WSN, mock-infected, or treated with hydrogen peroxide (H2O2; positive control). Cell lysates were blotted with reagents specific for total ADP-ribosylation (PAN), MARylation (MAR), or PARylation (PAR) modifications or the indicated proteins. * = unknown protein in infected cells migrating at ∼55 kDa, the molecular weight of influenza A virus NP. Lysates were probed for NP as a marker of infection and β-actin as a loading control. **(E)** Blotting of lysates from A549 cells infected or mock-infected with WSN with or without the PARG inhibitor PDD 00017273. Lysates were probed for NP as a marker of infection and β-actin as a loading control. **(F)** A549 cells were inoculated with infectious or UV-inactivated virus, or mock-treated. Total PARylation was detected in whole cell lysates by blotting. **(G)** Chicken UMNSAH/DF-1 fibroblasts, pig kidney epithelial PK(15) cells, and Brazilian free-tailed bat lung epithelial Tb 1 Lu cells were mock-infected or infected with WSN or a bat-adapted WSN for Tb 1 Lu cells. Total ADP-ribosylation and tubulin were detected by blotting. **(H-I)** Viral proteins are ADP-ribosylated. (H) Lysates from infected or mock-treated A549 cells were subject to immunoprecipitation with anti-NP of IgG control antibodies. Total PARylation was detected by blotting whole cell lysate (left) and immunoprecipitates (right). * = IgG heavy chain. (I) A549 cells were infected with WT virus encoding the FLAG-tagged polymerase subunit PB2. vRNPs were immunopurified from lysates with anti-FLAG antibody and blotted for total PARylation. Membranes were treated with hydroxylamine (HAM) where indicated prior to blotting. Images are from the same blot (Supplemental Figure 10). For B, data are presented as mean of 3 to 4 replicates ± SD. For C, data are grand mean of 2-3 biological replicates ± SEM. Multiple comparisons were made using a one-way ANOVA with *post-hoc* Dunnett’s test when compared to the control. * = p<0.05, ** = p<0.01, *** = p<0.001 and **** = p<0.0001. See also Figures S1-S3.

PARP8 was also identified with a correlation score similar to that of IFITM1-3 (Figure 1A). RNA-sequencing showed that PARPs are expressed in human lung A549 cells upon influenza virus infection or treatment with interferon β (IFNβ) (Figure S1A). Most PARPs are constitutively expressed, whereas several are specifically induced during the anti-viral response, including the known interferon-stimulated genes *PARP9, PARP10, PARP12, PARP13/Z3CHAV1*, and *PARP14* (Figure S1B)^10–14^.

Given the emerging role of PARPs at the virus:host interface^6–9,15^, we investigated the impact of PARP8 and other PARPs on influenza A virus replication. All tested PARPs, except PARP14, exhibited anti-viral activity upon over-expression in cells prior to infection, reducing titers for the influenza A virus strain A/WSN/1933 (H1N1; WSN) (Figure 1B). Validating the results of our screen, expression of PARP8 reduced viral gene expression and replication for diverse influenza A and B viruses (Figure S2A–C). To focus specifically on the effects of PARPs during genome replication and transcription, we measured polymerase activity in the presence or absence of PARP over-expression. Polymerase activity assays are performed by expressing the polymerase trimer, composed of subunits PB1, PB2 and PA, the viral nucleoprotein (NP), and a model genomic RNA encoding luciferase. These components assemble into ribonucleoprotein (RNP) complexes that replicate the viral genome and transcribe viral mRNA. Luciferase activity was measured to determine the cumulative output of these processes. Again, most PARPs exerted an anti-viral activity (Figure 1C). Our results agree with prior studies showing specific targeting of influenza A virus polymerase subunits by PARP13/Z3CHAV1^28,32^. Notably, PARP2, PARP8 and PARP10 did not affect the viral polymerase, but did interfere with viral replication, suggesting that they impact an polymerase-independent step in the viral life cycle. of ADP-ribosylation, resulting in robust detection of ADPr.

Because several PARPs were upregulated in response to FLUAV infection (Figure S1B), we investigated whether the enzymatic product of PARPs, ADP-ribosylation, was also upregulated. Basal levels of ADP-ribosylation were detected in mock-infected lung cells, whereas infection with influenza A virus increased total ADP-ribosylation, as stained by pan-ADPr reagent and antibodies specific for MAR or PAR (Figure 1D). The increase in MARylation is typified by the appearance of discrete ADP-ribosylated proteins on blots, while PARylation includes these bands, as well as higher-molecular weight smears. As a positive control, lung cells were also treated with hydrogen peroxide (H_2_O_2_), a known inducer ADP-ribosylation is highly dynamic and reversible through the activity of various enzymes such as poly(ADP-ribose) glycohydrolase (PARG)^24^. PARG cleaves ribosyl-ribosyl bonds in PAR chains. Therefore, we repeated infections in the presence of a PARG inhibitor (PDD 00017273). PDD treatment increased the PARylation signal from infected cells compared to untreated infected cells, whereas ADP-ribosylation was largely unchanged in all conditions for mock-infected cells (Figure 1E). These data indicate that infection causes a marked induction of PARylation. Thus, PDD treatment was included in subsequent experiments to facilitate detection of ADP-ribosylated proteins. Next, we exposed cells with infectious virus, or the same dose of UV-inactivated particles. Only cells exposed to replication-competent virus showed an increase in ADP-ribosylation (Figure 1F). Viral exposure also up-regulated ADP-ribosylation in cells derived from chicken, pig and bats (Figure 1G), all of which are natural hosts of influenza A virus^33^. Thus, global ADP-ribosylation changes are a common response to active replication of influenza A virus.

To specifically test whether infection-induced ADP-ribosylation results in modification of viral proteins, we immunopurified NP from infected cells (capturing NP, as well as NP assembled into RNPs with the viral polymerase). Blotting of NP purified from infected cells revealed robust ADP-ribosylation indicative of PARylation, which was absent in control conditions (Figure 1H).

In an orthogonal approach, we first enriched ADP-ribosylated proteins from cell lysates using the recombinant macrodomain protein Af1521, which has high affinity for ADPr, and then probed for NP. NP was selectively captured by Af1521, but not Af1521 G42E, an ADPr-binding mutant (Figure S3A). To further confirm the specificity of blotting and ensure detection of *bona fide* ADP-ribosylation events, we immuno-purified RNPs from infected cells and split samples into parallel blots. For the control membrane, western blots revealed ADP-ribosylation consistent with modifications to NP and the viral polymerase (Figure 1I, left). The other membrane was treated with hydroxylamine (HAM) (Figure 1I, right), which releases ADPr conjugates, primarily from modified aspartyls, glutamyls and arginyls^34^. HAM treatment markedly reduced detection of ADP-ribosylation, confirming that viral proteins are direct targets of PARP activity. HAM treatment did not completely eliminate the signal; fainter signal was still detected at the expected molecular weight of NP, suggesting that it may be modified at residues resistant to HAM treatment (e.g. histidyls, cysteinyls, seryls or threonyls). Together, these data show that influenza A virus infection induces expression of multiple PARPs and causes a global up-regulation of ADP-ribosylation, including the modification of viral proteins.

### Global identification of ADP-ribosylation sites during infection includes functionally important modification of viral proteins

Despite the broad impact of ADP-ribosylation on viral infections^7^, global analyses of ADP-ribosylated proteins have largely been studied in the context of DNA damage and oxidative stress^35,36^. Therefore, we evaluated how influenza A virus infection alters the ADP-ribosylated proteome using enzymatic labeling of terminal ADPr-coupled mass spectrometry (ELTA-MS)^36,37^. ELTA selectively modifies ADP-ribosylated peptides with “clickable” *N*-azido-labeled tags, thereby enabling enrichment using click chemistry on a solid support, followed by stringent washes prior to MS for site identification. We identified the ADP-ribosylome in two biological replicates of A549 lung cells infected with the WSN strain of influenza A virus, mock-infected cells, or treated with hydrogen peroxide (H_2_O_2;_ positive control). Each sample was analyzed in technical triplicate. A peptide counted as ADP-ribosylated when it was detected in at least two technical replicates, one of which was by direct identification via tandem mass spectrometry (MS/MS). There was a high degree of overlap between technical and biological replicates, validating our approach (Figure S4A, Table S1). In total, across all conditions, even with these very strict cutoffs, we identified ∼1,200-1,300 ADP-ribosylation sites on 600–700 proteins (Figure 2A–B).

**Figure 2.**
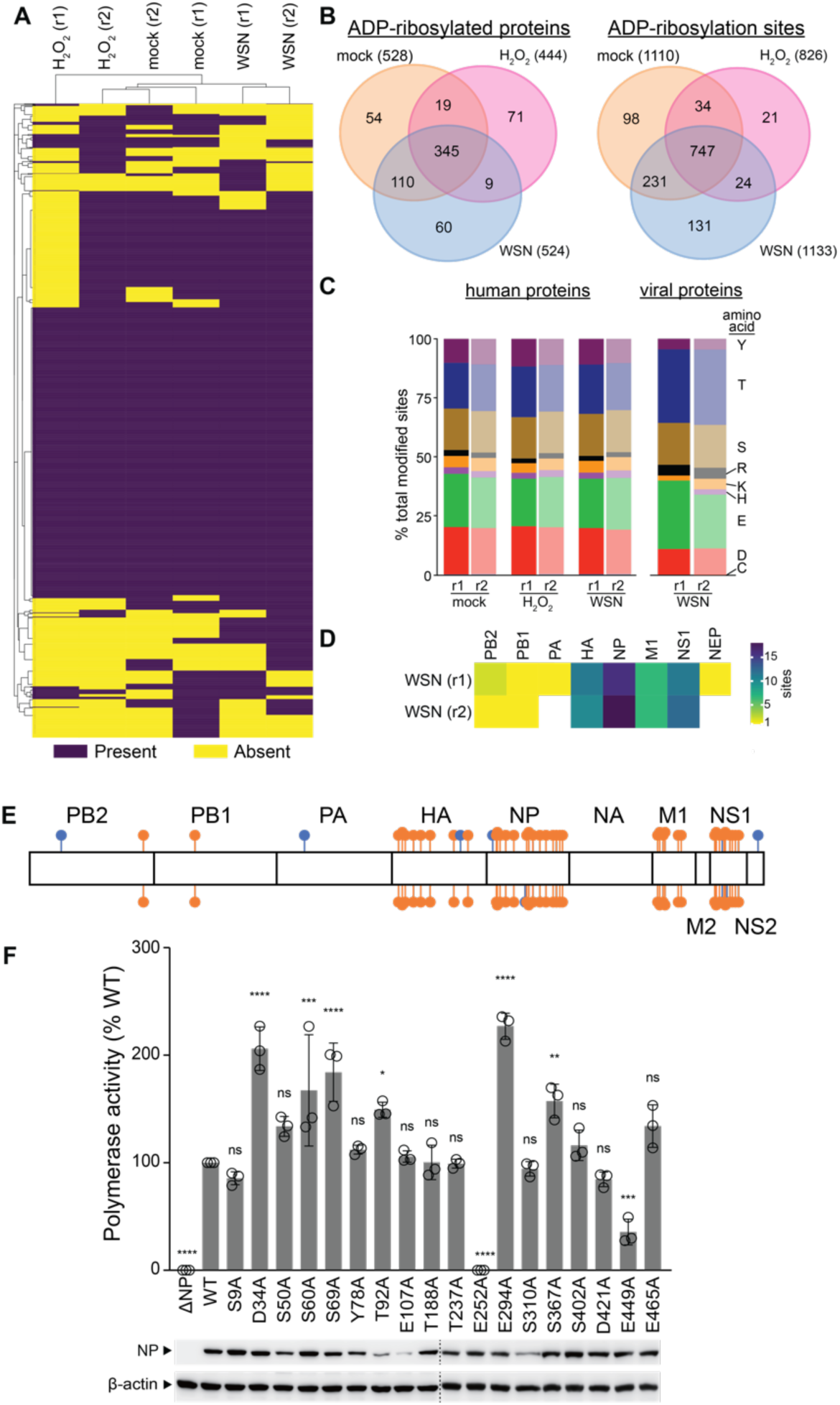
Identification of ADP-ribosylation sites on viral and host proteins. Human lung A549 cells were infected with WSN, mock infected, or treated with H2O2. ADP-ribosylation sites were identified by ELTA-MS. **(A)** Heat map clustering unique ADP-ribosylation sites of human and viral proteins in two biological replicates. **(B)** Venn diagrams of modified proteins (top) and sites (bottom) identified in (A). **(C)** Distribution of ADP-ribosyltions detected on the specified amino acid residues of human (*left)* or viral (*right*) proteins. Note that there were no modified cysteines in viral proteins. **(D)** Heatmap of the number of ADP-ribosylation sites identified on viral proteins during infection. **(E-F)** ADP-ribosylation sites are important for influenza A virus protein function. (E) ADP-ribosylation sites mapped on to a concatenated influenza A virus proteome. Sites identified in both biological replicates are marked in orange, whereas those unique to a replicate are in blue. (F) Ablation of ADP-ribosylation sites alters NP function. Polymerase activity assays performed in 293T cells with WT NP or ADP-ribosylation mutants. Expression of NP and mutants was confirmed by immunoblotting, along with β-actin as a loading control. For each condition, ELTA-MS was performed in technical triplicate on two independent biological replicates. Data in F are mean of n=3 ± SD. * = p<0.05, ** = p<0.01, *** = p<0.001 and **** = p<0.0001 as determined by a one-way ANOVA with Dunnett’s multiple comparisons test when compared to WT. See also Figures S3-S4 and Table S1.

While infection produces a distinct ADP-ribosylation profile in A549 cells, there was a high degree of overlap among experimental conditions (i.e., mock, infected and H_2_O_2_-treated). Over half of the modified proteins, and almost three-quarters of all modified sites, were shared across all three conditions with a similar distribution of amino acid residues that were modified (Figure 2B–C). This collection of shared proteins was highly enriched in gene ontology (GO) terms for protein translation (Figure S4B).

We also identified ADP-ribosylation sites on eight of the ten canonical influenza A virus proteins (Figure 2D–E). The distribution of amino acids modified in viral proteins was similar to that of host proteins, with the exception that there were no modified cysteinyls detected in viral proteins (Figure 2C). NP repeatedly had the highest number of modifications, consistent with our biochemical approaches that detected ADP-ribosylated NP (Figure 1H–I). It is unclear whether this is because NP is the most frequently modified viral protein, or because it is one of the most abundant proteins in infected cells. Together, these findings suggest the presence of a core ADP-ribosylome in A549 cells, with different stimuli both increasing and decreasing modifications on distinct subsets of proteins, and show that influenza A virus induces a unique profile of ADP-ribosylated proteins and modification sites, including viral proteins.

Notably, although blotting revealed massive increases in PARylation, ELTA-MS showed that the total number of modified proteins or sites does not change (Figure 1D–H versus Figure 2A–B). ELTA-MS uses a phosphodiesterase to cleave the pyrophosphate bond in ADPr to leave a single phosphoribosyl moiety that enables an unambiguous identification of ADP-ribosylation sites^36^. However, this cleavage eliminates the length information originally associated with the modified site (e.g., MAR versus PAR). The apparent contradiction between our blotting and MS suggests that infection does not alter which sites are modified, but instead changes the ADP-ribosylation form from MAR to PAR.

A panel of 18 NP variants representing high-confidence modification sites was used to determine the functional effects of ADP-ribosylation (Table S1). ADP-ribosylation sites were found in the head and body domains of NP, the tail loop that mediates NP oligomerization, and flanking the positively-charged RNA binding groove (Figure S3B–C). ADP-ribosylation sites were ablated by changing the modified residues to alanyls, and NP function was assessed in a polymerase activity assay. Consistent with the finding that PARP expression inhibited FluPol activity (Figure 1C), polymerase activity increased for several of the NP ADPr-site variants, with the most pronounced effects observed for variants D34A, S69A, and E294A (Figure 2F). In some cases (e.g. NP T92A E107A), variants resulted in activities approaching those in control conditions with unchanged NP even though the amino acid substitutions appeared to affect the overall stability or expression of NP, perhaps masking a net increase in NP activity (Figure 2F). Curiously, the E252A variant resulted in a complete loss of polymerase activity (Figure 2F), in agreement with prior work^38^. Conversely, deep mutational scanning of NP selected for changes at residue 294 to amino acids that cannot be ADP-ribosylated, reinforcing that ADP-ribosylation may attenuate NP function (Figure 2F)^38^.

### Influenza A virus NS1 suppresses global ADP-ribosylation

Our data indicated that ADP-ribosylation is part of the anti-viral response to influenza A virus infection. Influenza virus NS1 is the primary viral protein that antagonizes host anti-viral responses and was previously shown to reduce ADP-ribosylation of argonaute RISC catalytic component 2 (AGO2) complexes^39,40^. Therefore, we investigated the impact of NS1 on global ADP-ribosylation by infecting cells with two closely related strains of influenza A virus encoding NS1 proteins of different potency: WSN, which was used in experiments described above, and A/Puerto Rico/8/1934 (H1N1; PR8). Both strains induced ADP-ribosylation (Figure 3A). Although global ADP-ribosylation was lower in cells infected with PR8 compared to WSN, despite similar infection, levels, PARylation was transiently induced regardless of the strain, as indicated by the ability of PDD to preserve the signal. PARP1, in addition to functioning as a PARP, is a target for caspases and its cleavage is frequently used as a marker for the activation of apoptosis^41,42^. Apoptosis plays dual pro- and anti-viral roles during infection, and its regulation by influenza A virus is important for successful replication^43–45^. In addition to higher levels of PARylation, PARP1 cleavage was also more pronounced in cells infected with WSN compared to PR8 (Figure 3A).

**Figure 3.**
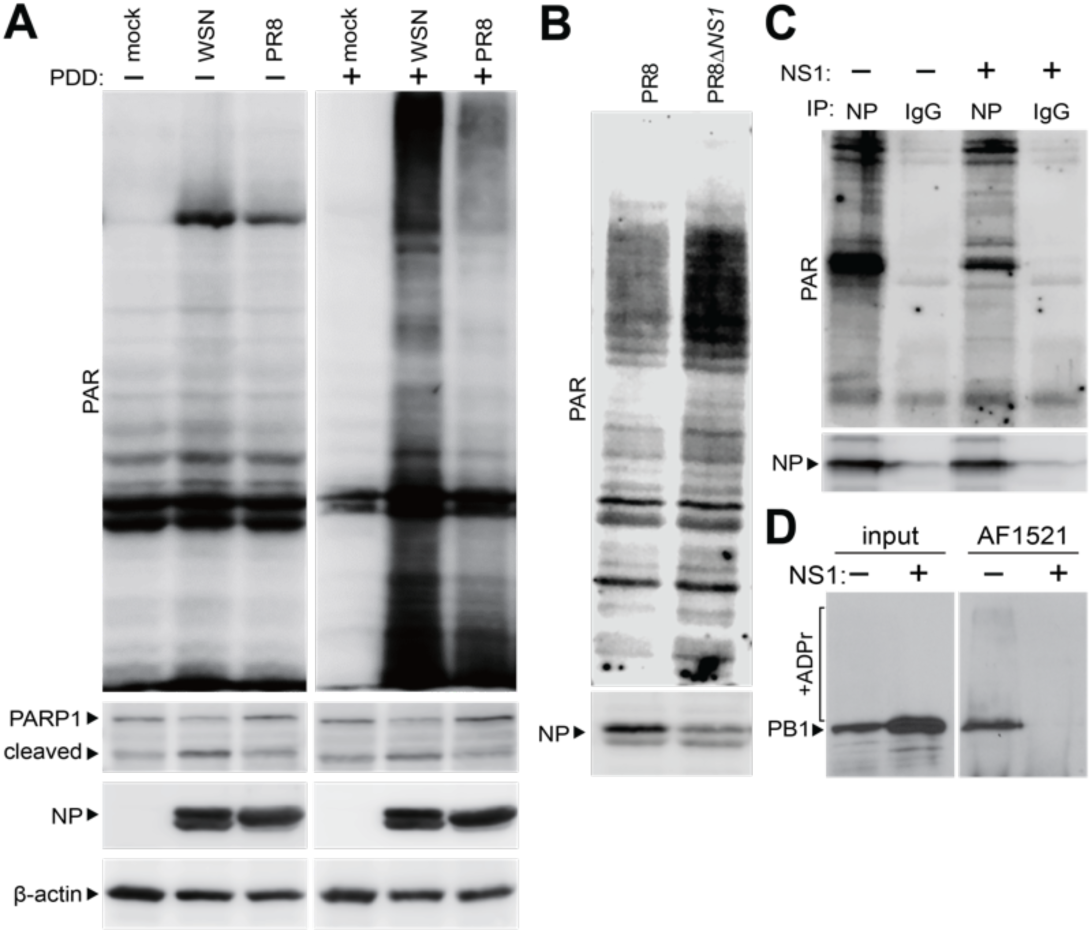
Induction of ADP-ribosylation differs across closely related strains of influenza A virus. **(A)** Strain-specific differences in ADP-ribosylation. Human lung A549 cells were infected with WSN or PR8, or mock infected, and treated with the PARG inhibitor PDD 00017273 where indicated. Proteins and ADP-ribosylation were detected by blotting. Lysates were probed for NP as a marker of infection and β-actin as a loading control. **(B)** NS1 counters ADP-ribosylation. ADP-ribosylation was detected in whole cell lysates from A549 cells infected with PR8 or PR8 lacking NS1 (PR8Δ*NS1*). Lysates were probed for NP as a marker of infection. **(C-D)** Influenza virus NS1 protein suppresses ADP-ribosylation of viral proteins. Influenza A virus RNPs were assembled in 293T cells in the presence or absence of NS1. (C) RNPs were immunopurified with anti-NP antibody and probed for ADP-ribosylation or NP. (D) ADP-ribosylated proteins were affinity purified with recombinant Af1521. The influenza polymerase subunit PB1 was detected in the input and purified fractions by blotting.

Next, we tested a direct role of NS1 in regulating PARylation by infecting cells with a virus that lacks the *NS1* gene (PR8Δ*NS1*). PARylation was dramatically increased in cells infected with PR8Δ*NS1* compared to PR8, even though PR8Δ*NS1* virus is highly attenuated as revealed by lower levels of NP (Figure 3B). NS1 expression also reduced viral protein PARylation. Influenza A virus RNPs were assembled by ectopic expression in cells in the presence or absence of NS1. RNPs were immunopurified with antibodies against NP and probed for ADP-ribosylation, revealing much less PARylation when NS1 was present (Figure 3C). In a reciprocal experiment, we enriched ADP-ribosylated proteins using recombinant Af1521 and blotted for the viral polymerase subunit PB1. PB1 was captured by Af1521, indicative of ADP-ribosylation, but no capture occurred when NS1 was co-expressed (Figure 3D). Combined, these data show that NS1 antagonizes global PARylation during infection while also reducing modification of viral proteins.

To further investigate the kinetics of PARylation and the associated role of NS1, we performed single-cycle infections in A549 cells with PR8 or PR8Δ*NS1*. For PR8, increased PARylation was detected at 8 hpi, i.e. in a later stage of infection (Figure 4A). PARylation levels then continued to increase over the time course. In contrast, PARylation was apparent as early as 4 hpi in cells inoculated with PR8Δ*NS1*. As before, PARylation continued to increase over the time course, but at much higher levels. The increase in PARylation was associated with higher levels of PARP1 cleavage (Figure 4A). NP is also a target for caspase cleavage, resulting in the removal of ∼16 amino acids from the N-terminus^46^. Blotting revealed earlier NP cleavage in PR8Δ*NS1*-infected cells compared to PR8-infected cells, again linking increased PARylation with apoptotic activity (Figure 4A).

**Figure 4.**
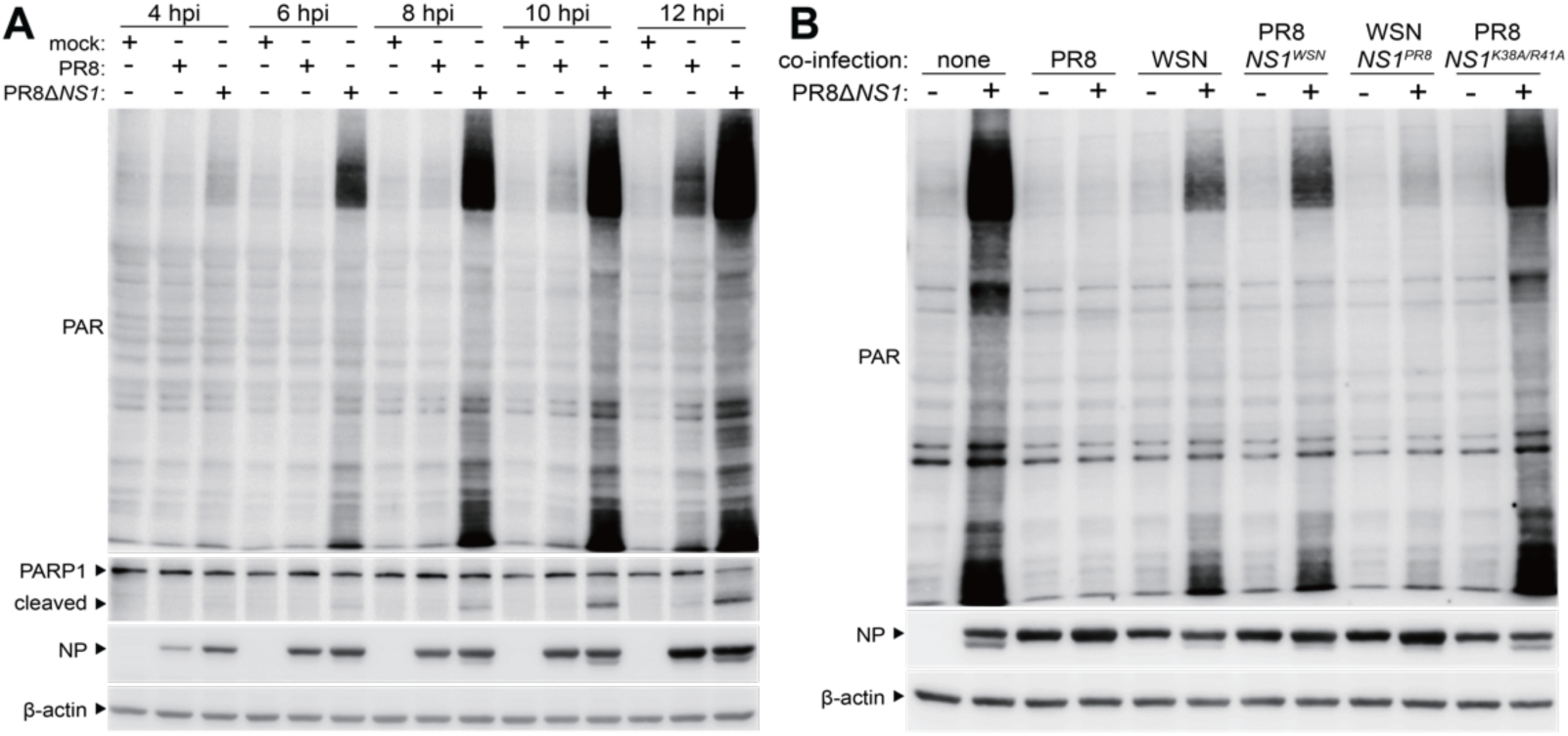
Influenza A virus NS1 tempers ADP-ribosylation by the host. **(A)** NS1 suppresses infection-induced ADP-ribosylation throughout infection. Proteins and global ADP-ribosylation were detected by blotting lysates from human lung A549 cells infected with PR8 (MOI=5) or PR8Δ*NS1* (MOI=5) or mock infected. **(B)** Virus-specific differences in ADP-ribosylation map to NS1 and require its RNA-binding activity. A549 cells were infected with PR8Δ*NS1* (MOI=4.5) and co-infected with virus encoding the indicated heterotypic NS1 or the RNA-binding mutant NS1 K38A/R41A (MOI=0.5). Infections proceeded for 8 hr prior to lysis and blotting.

We capitalized on the robust ADP-ribosylation during PR8Δ*NS1* infection to test functions of NS1 that are important for controlling PARP activity. We induced PARylation by inoculating cells with PR8Δ*NS1* and then attempted to rescue the Δ*NS1* phenotype by providing NS1 in *trans* by co-infection with virus encoding different NS1 proteins. Paralleling prior results, PR8 NS1 completely suppressed PARylation while WSN NS1 reduced, but did not eliminate it (Figure 4B). We confirmed that this difference between PR8 and WSN was due solely to NS1 by swapping this gene between each virus. WSN encoding *NS1^PR^*^8^ (WSN *NS1^PR^*^8^) was now able to potently suppress PARylation, whereas PR8 encoding *NS1^WSN^* (PR8 *NS1^WSN^*) caused less suppression (Figure 4B). One of the major mechanisms by which NS1 antagonizes host responses is through binding double-stranded RNA to prevent its sensing by innate immune activators^47,48^. We therefore generated virus encoding a variant that no longer binds double-stranded RNA (NS1 R38A/K41A)^47^. NS1 R38A/K41A lost its ability to suppress PARylation, with results indistinguishable from infection by PR8Δ*NS1* alone (Figure 4B). These data reveal that NS1 counteracts the anti-viral activity of PARPs and ADP-ribosylation, at least in part through its ability to bind dsRNA.

### NS1 suppresses PARylation but does not reduce the number of ADP-ribosylation sites

NS1 suppresses global PARylation, leading us to hypothesize that NS1 controls the number of sites that are modified. Infection with a virus lacking NS1 would thus be expected to increase the number of ADP-ribosylation sites. We tested this possibility by again performing ELTA-MS, this time comparing infections in the presence or absence of NS1. In one set of experiments, we compared infections with WSN, PR8, and PR8ΔNS1 (ELTA-MS #2), and in the other set we also included WSNΔ*NS1* (ELTA-MS #3), using the same controls as before. Because viruses encoding NS1 deletions are attenuated, infection conditions were optimized for each virus to ensure equivalent expression levels of viral proteins. As before, we identified ∼ 2,600-3,000 modified sites in ∼380-660 proteins (Figure 5A-B, Figure S5, Table S2-S3).

**Figure 5.**
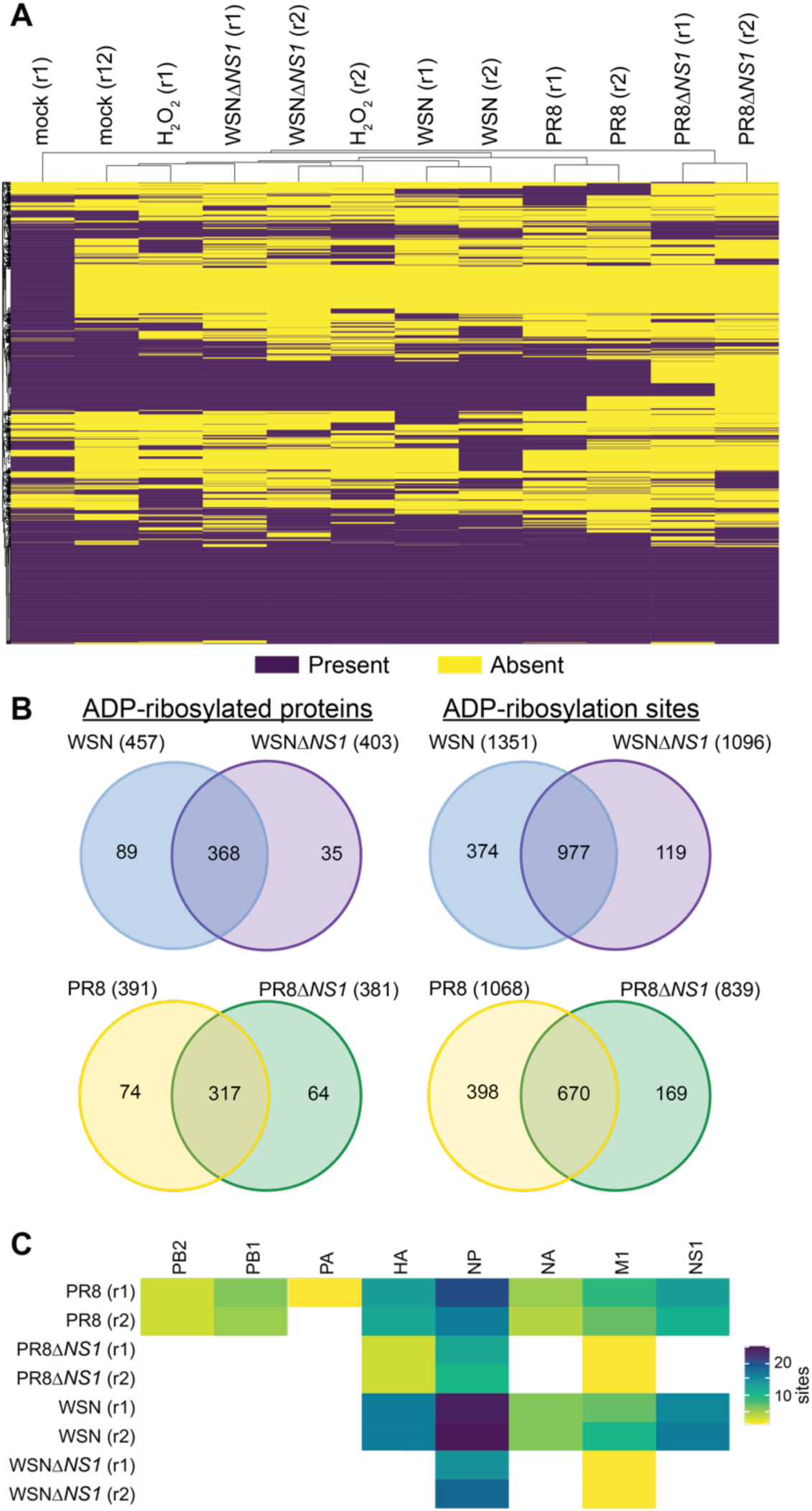
ADP-ribosylation in the absence of NS1. Human lung A549 cells were infected with PR8, PR8Δ*NS1*, WSN,WSNΔ*NS1*, mock infected, or treated with H2O2. ADP-ribosylation sites were identified by ELTA-MS. **(A)** Heat map clustering unique ADP-ribosylation sites of human and viral proteins present in two biological replicates. **(B)** Venn diagram of the number of modified proteins (left) and sites (right) identified (A). **(C)** Heatmap of the number of ADP-ribosylation sites identified on viral proteins during infection. For each condition, ELTA-MS was performed in technical triplicate on two independent biological replicates. See also Figure S5-6 and Tables S2-3.

Across all datasets, we detected ADP-ribosylation of proteins involved in diverse cellular functions, including consistent enrichment in proteins involved in protein binding, RNA binding, cadherin binding and cell adhesion molecule binding (Table S4). Infections induced a unique profile of ADP-ribosylation, while significant overlap among conditions reinforced the idea of a core ADP-ribosylome in A549 cells. Most of the modifications were shared across independent experiments, highlighting the reproducibility of ELTA-MS (Figure S6A). There was a high degree of overlap among modified proteins and sites for cells infected with WSN or PR8 (Figure S6B). Remarkably, despite large differences in the intensity of PARylation revealed by blotting, cells infected with WT and Δ*NS1* mutant virus had comparable numbers of ADP-ribosylation sites and modified proteins, with the majority of sites and proteins found in both conditions (Figure 5B, Figure S5B). This result was consistent among WSN and PR8 strains. ADP-ribosylation sites were identified on up to eight different viral proteins (Figure 5C, Figure S5C). Regardless of the presence or absence of NS1, NP was the viral protein with the greatest number of ADP-ribosylation sites. Contrary to our original hypothesis, these data show that the number of sites identified during infection did not increase when *NS1* was deleted or when comparing the weak NS1 expressed by WSN against the more potent NS1 from PR8. If anything, infection with Δ*NS1* viruses resulted in detection of slightly fewer ADP-ribosylated proteins and ∼20% fewer sites. Therefore, NS1 likely suppresses PARylation not by reducing the number of sites, but by reducing the number of ADPr units at each site by inhibiting the transition from MARylation to PARylation.

### PARP1 directs anti-viral PARylation

Our data suggest that infection primarily changes PARylation. PARylation is performed by a limited subset of PARPs – PARP1, PARP2, PARP5a (TNKS) and PARP5b (TNKS2)^18^. ELTA-MS identified ADP-ribosylation at two canonical auto-modification sites in PARP1, E488 and E491, indicating that at least PARP1 was active in our infected cells (Table S2-S3)^49^. To specifically test which PARPs are involved in the response to influenza virus infection, we measured ADP-ribosylation in infected cells treated with a series of inhibitors possessing different specificities: PJ34 generically inhibits PARPs; Olaparib and Rucaparib broadly target PARP1 and PARP2, although they have been shown to possess off-target activity against other PARPs; AG14361 selectively inhibits PARP1; and, XAV939 primarily targets PARP5a and PARP5b^17,50^. ADP-ribosylation was unaffected by XAV939 in cells infected with either WSN or PR8, eliminating PARP5a/TNK and PARP5b/TNKS2 as candidates (Figure 6A). By contrast, PARylation was inhibited by Olaparib, Rucaparib, and the PARP1-specific AG14361, implicating PARP1 as the dominant enzyme mediating PARylation. Interestingly, caspase cleavage of NP still occurred under all conditions, indicating that infection-induced apoptosis does not require PARylation (Figure 6A). Next, we purified viral RNPs from infected cells and probed them for MARylation or PARylation. Treating infected cells with Olaparib did not affect MARylation of the purified RNPs, but dramatically reduced the degree of PARylation (Figure 6B). We generated two independent *PARP1* knockout A549 cell lines to directly test its activity during infection (Figure 6C). We infected these cells with PR8ΔNS1 to create a highly sensitized setting to observe changes in ADP-ribosylation. Infection-induced PARylation was almost completely absent in both *PARP1* knockout cell lines (Figure 6D). We repeated these experiments using WT PR8 and WSN. Again, the global increase was absent in *PARP1* knockouts (Figure 6E). Together, these data show that PARP1 mediates the majority of PARylation on viral proteins as well as a global increase in the cell, whereas viral proteins are MARylated by an unknown PARP.

**Figure 6.**
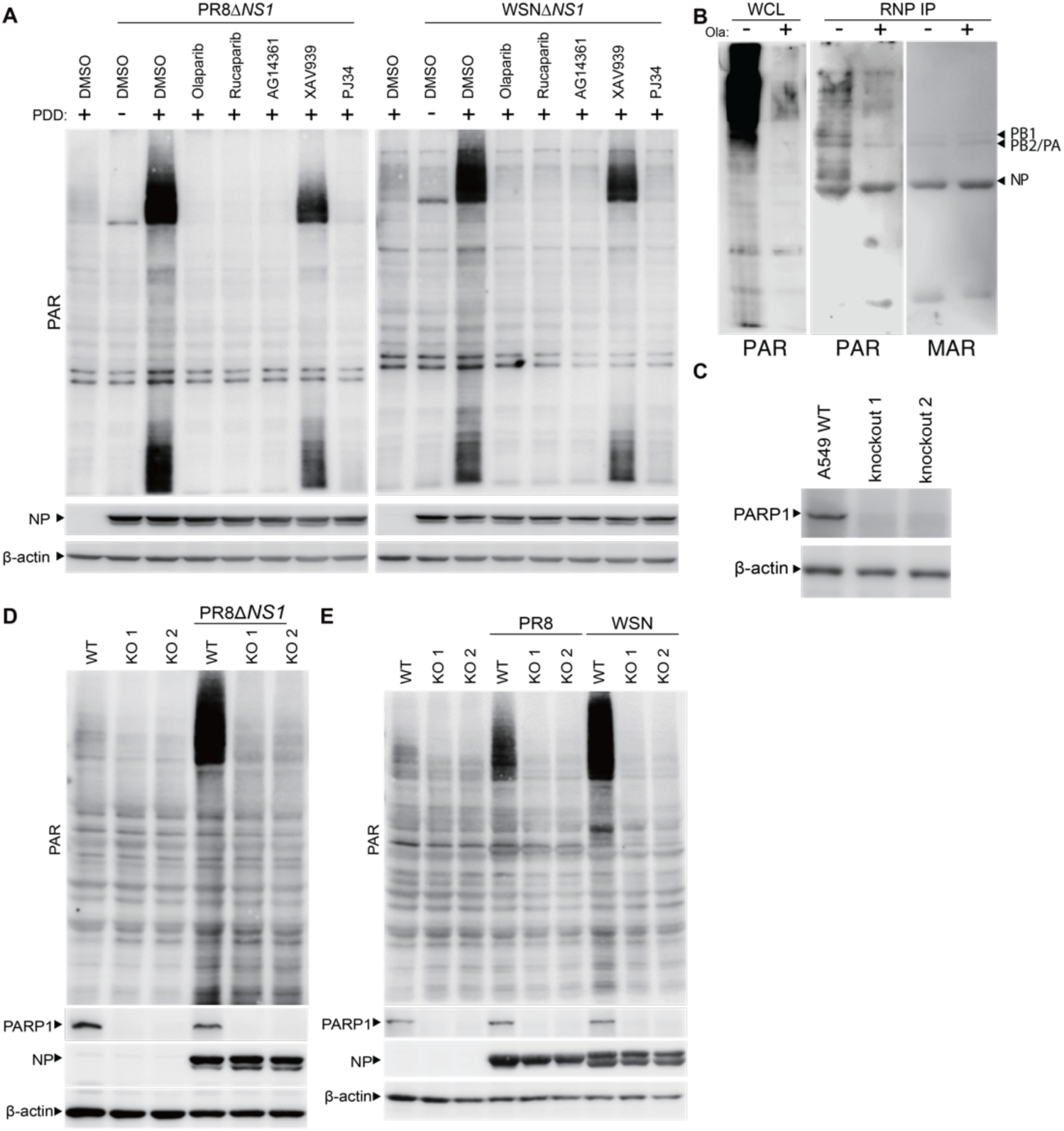
PARP1 regulates PARylation during influenza A virus infection. **(A)** Chemical inhibitors identify PARP1 as the primary PARP active during influenza A virus infection. ADP-ribosylation was measured by blotting lysates from mock-treated or infected human lung A549 cells treated with the indicator PARP inhibitors or a DMSO control. **(B)** PARP1 modifies viral proteins. A549 cells were infected with WSN PB2-FLAG and treated with Olaparib or DMSO. Influenza A virus RNPs were immunopurified from infected cell lysate. Samples were probed with reagents specific for PAR or MAR. **(C)** Validation of PARP1 knockout in two clonal A549 cell lines by western blotting. β-actin was targeted as a loading control. **(D-E)** PARP1 mediates ADP-ribosylation in IAV-infected cells. WT A549 or PARP1 knockout (KO) lines were infected with (D) PR8Δ*NS1* or (E) WT PR8 and WSN. ADP-ribosylation and proteins were detected by blotting whole-cell lysate. Lysates were probed for NP as a marker of infection and β-actin as a loading control.

PARP1 both initiates ADP-riboyslation via MARylation and extends chains via PARylation^51^. We investigated these activities by complementing our knockout cells with PARP1, or PARP1 E988A (inactive catalytic site), or PARP1 E988K (only able to MARylate)^51,52^. Complementation with WT PARP1 restored global ADP-ribosylation and PARylation in cells infected with WSN or PR8 as revealed by blotting for PAR and pan-ADPr (Figure 7A, Figure S7A). This result confirms that the knockout cell phenotype was due to the loss of PARP1, rather than off-target effects. As expected, the PARP1 variants did not restore PARylation. However, neither PARP1 E998A nor PARP1 E998K caused global changes in MARylation, with levels similar to the knockout cells. There was an observable decrease in MARylation in A549 *PARP1* knockout cells compared to the parental cells, overlapping with the high-molecular smearing in the PAR blot. Whether this represents actual MARylation by PARP1, or results from the low levels of cross-reactivity reported for the MAR detection reagent, needs to be determined^53^. Curiously, complementation with a PARP1 variant D214A, which cannot undergo caspase cleavage, failed to restore PARylation in infected cells (Figure S7B). These data indicate that PARP1 drives global PARylation during infection, but that other PARPs—including the MAR-adding PARP8 found in our initial screen (Figure 1A)—likely contribute the majority of MARylation.

**Figure 7.**
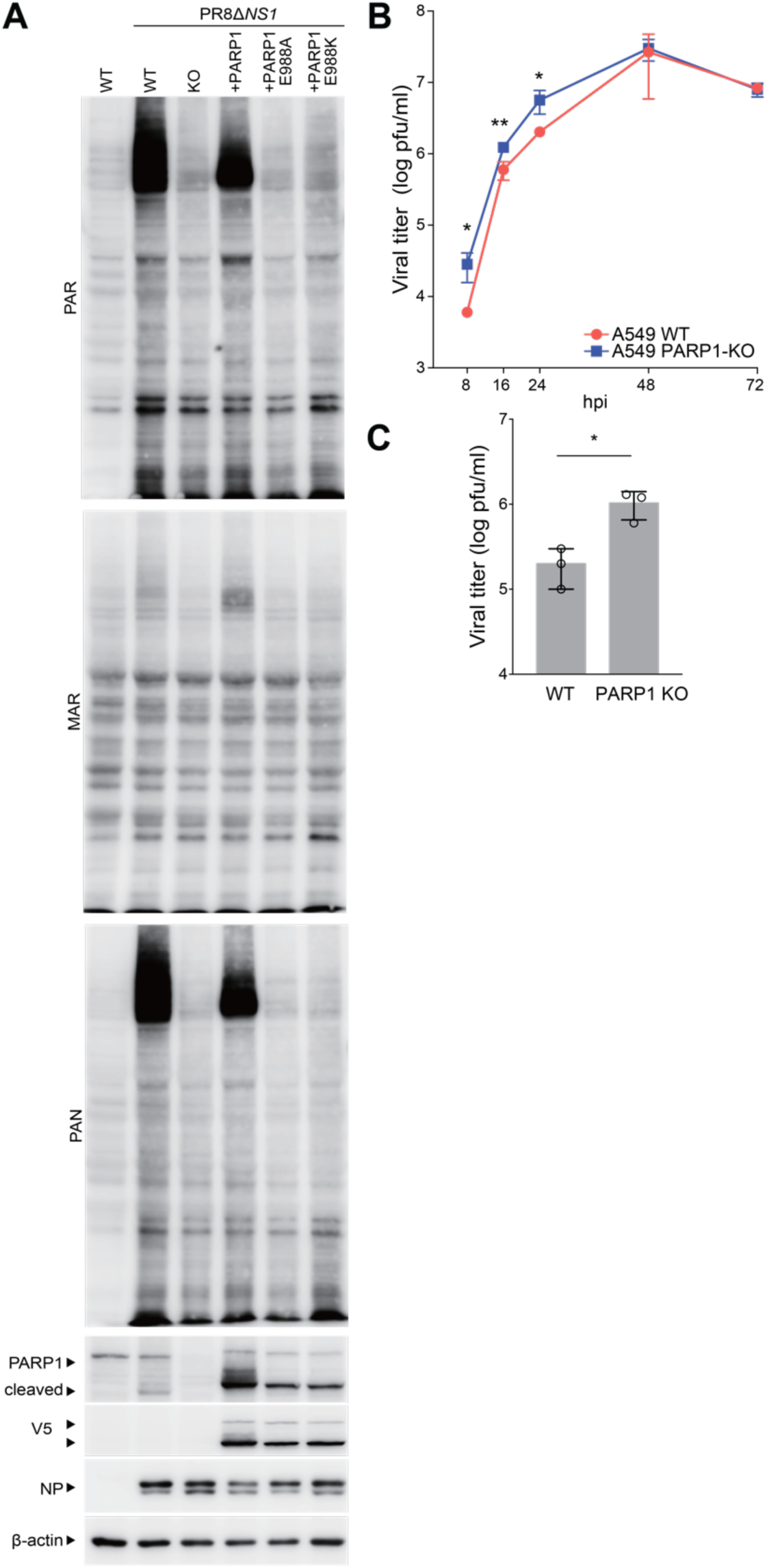
PARP1-mediated ADP-ribosylation suppresses influenza A virus replication. **(A)** PARylating activity of PARP1 is required for infection-induced ADP-ribosylation. Human lung A549 *PARP1* knockout cells were complemented with PARP1-V5 or the PARylating mutants E988A or E988K. Cells were infected or mock treated and whole cell extracts were used for blotting. Lysates were probed for NP as a marker of infection and β-actin as a loading control. **(B-C)** PARP1 suppresses influenza A virus replication. (B) Multi-cycle and (C) single-cycle replication of PR8 in parental A549 cells or A549 *PARP1* knockout cells. Viral titers were determined by plaque assay for the timepoints indicated in the multicycle assay or 8 hpi for the single-cycle assay. Data are mean of n=3 ± SD. * = p<0.05 and ** = p<0.01 as determined by t-test when compared to WT. See also Figure S7.

Finally, we tested the effect of PARP1 on viral replication by infecting *PARP1* knockout cells. Multi-cycle replication assays were initiated in A549 and *PARP1* knockout lung cells. Virus replicated to ∼five-fold higher titers in cells lacking PARP1 during the early stages of infection (Figure 7B). Viral titers ultimately reached the same plateau in both cell lines. In a single-cycle replication assay, *PARP1* knockout cells again produced five-fold more virus than paretnal A549 cells (Figure 7C). Thus, PARP1 orchestrates a global increase in PARylation during infection, establishing an anti-viral activity that slows the replication of influenza A virus.

## DISCUSSION

Viral infections trigger innate immune responses that direct cell-autonomous anti-viral responses. Here we show that influenza A virus infection induces PARylation that antagonizes viral replication. Host and viral proteins are modified as part of an infection-specific pattern. This was characterized not by a global change in the proteins that were ADP-ribosylated, but rather by a transition from MARylation to PARylation for those that were already modified. Polymerase activity assays showed that specific modifications on viral proteins disrupt their function, suggesting that ADP-ribosylation directly impacts the viral replication machinery. The degree of ADP-ribosylation was strain-dependent, with the differences mapping to the viral NS1 protein. NS1 was shown to suppress ADP-ribosylation, primarily by preventing formation of PAR chains. PARP1 was identified as the cellular enzyme responsible for this PARylation and conferring the anti-viral phenotype. These data show that ADP-ribosylation is an important component of the anti-viral response to influenza A virus.

Our results reveal a core ADP-ribosylome that is modified to create an infection-specific profile. Across all of our infections, we identified 4,064 unique modification sites on 1,002 proteins. Although infection causes new modifications on a small number of proteins, most changes appear to involve PARylation at existing modification sites. The smaller proteome of influenza A virus has enabled us to define the functional consequences of ADP-ribosylation at specific sites on viral proteins. A total of 27 ADP-ribosylation sites were identified in NP across all of our replicates, including 18 high-confidence sites that we investigated in polymerase activity assays (Figure 2 and 5, S3B, S8, S9, Tables S1-3). NP binds RNA and assembles into homo-oligomers along the length of the viral genome, forming the scaffold on which viral transcription and replication take place^54^. NP binds RNA in a sequence-independent fashion using a large basic face to interact with the phosphate backbone^55–57^. Recent crystal structures identified specific residues in NP that contact the RNA, including S69, T92, and S367^55^, which we have also identified as high-confidence ADP-ribosylation sites. Mutating these residues to prevent ADP-ribosylation enhanced polymerase activity, indicating that ADP-ribosylation at these positions disrupts function, possibly by precluding binding to the viral genome. This finding also raises the intriguing possibility that the negatively charged phosphate backbone of longer ADPr chains may compete for binding not just at the residues that are modified, but across the larger positively charged RNA-binding surface of NP. Thus, at these sites, ADP-ribosylation, especially the polymeric form, may mimic nucleic acids to disrupt assembly of the replication machinery.

Different forms of ADP-ribosylation are critical for directing cellular functions. *PARP1* knockout cells are deficient in DNA repair^58^. Wild-type, PARylation-competent PARP1 is able to fully rescue DNA repair in *PARP1* knockout cells, but PARP1 E988K, which is only capable of MARylation, cannot^59^. We observed a significant induction of PARylation during influenza infection, independent of virus strain, which is dependent on PARylation-competent PARP1. PARP1 confered anti-viral activity during influenza A virus infection, yet the PARP family possesses dual pro- and anti-viral functions. PARP1, PARP7, and PARP11 have each been reported to increase viral replication by reducing type I interferon production or signaling^60–62^. Conversely, PARP7 and PARP11, along with PARP1, PARP9, PARP10, PARP12 and PARP13/Z3CHAV1 all antagonize infection^6,7^. It was recently suggested that PARP11 antagonizes interferon responses^61^, but its expression in our studies clearly showed anti-viral activity against influenza A virus. PARP14 also possesses dual pro- and anti-viral activity, depending on the virus under investigation^63^. An anti-viral phenotype is not always dependent on catalytic activity, as PARP13/ZC3HAV1 exerts broad-spectrum anti-viral activity despite being catalytically inactive^7^. Both the long and short isoforms of PARP13/ZC3HAV1 inhibit influenza A virus replication. PARP13L coordinates PARylation of the viral polymerase by an unknown PARP(s) and subsequent ubiquitin-mediated degradation^28^. As both PARP1 and PARP13/Z3CHAV1 interact with the viral polymerase, it is possible that that anti-viral activity of PARP1 we defined here may be contributing the ADP-ribosylation coordinated by PARP13L^26,28^. This contrasts with PARP13S, whose anti-viral activity appears to be independent of ADP-ribosylation^29^. Thus, the ultimate effect of PARPs and ADP-ribosylation during infection is likely contextual, depending on the virus, the cell, and the host.

Given the inhibitory effects of specific PARPs on viral replication, it is unsurprising that viruses have evolved mechanisms by which they combat PARPs and ADP-ribosylation. Diverse positive-sense RNA viruses including alphaviruses, coronavirids, rubella virus, and hepatitis E virus encode macrodomain proteins that hydrolyze and thereby remove ADP-ribosylation modifications from proteins. Viral macrodomains are necessary for successful replication^6,8^. Indeed, small molecule inhibitors of the severe acute respiratory syndrome coronavirus 2 (SARS-CoV-2) macrodomain Mac1 inhibit viral replication^64^. The influenza A virus genome does not encode a macrodomain, but it does encode the potent immune antagonist NS1^65^. NS1 is not known to bind ADPr or possess ADP-ribosylhydrolase activity. Nonetheless, NS1 suppresses global PARylation as well as modification of specific proteins, with the potency of this activity varying between strains (Figure 3-4)^40^. Our data show that NS1 RNA-binding mutants no longer suppress ADP-ribosylation. This finding agrees with prior work using an NS1 phosphomimetic that prevents RNA binding^40^. As the RNA binding activity of NS1 has pleiotropic effects, it is not clear if the antagonism of ADP-ribosylation is direct or indirect. Influenza A virus appears to possess a unique strategy to combat PARPs and ADP-ribosylation.

The molecular signatures that activate PARPs during influenza A virus infection are unknown, although the data with NS1 raise the possibility that viral dsRNA may broadly be a trigger for global PARylation. This model is supported by the observation that many of the inhibitory effects of PARPs have been established in the context of RNA viruses, suggesting that PARP family members are players in the anti-viral RNA-sensing pathway, and hence could be activated indirectly by known RNA sensors, such as RIG-I, MDA5 or PKR^66^. However, it is unlikely that RIG-I is a major player in this process because global ADP-ribosylation was detected in chicken cells, which lack RIG-I (Figure 1G)^67^. A more provocative hypothesis is that PARPs directly sense foreign RNAs^68^. PARP1 is activated by binding cellular small nucleolar RNAs (snoRNAs)^69,70^. Whether viral RNAs can trigger the same response, or even host RNAs that are expressed in response to infection, remains to be determined. In summary, we established PARPs and ADP-ribosylation as new players in the battle against influenza A virus and highlight how the virus parries this attack.

## ACKNOWLEDGEMENTS

We thank B. tenOever for reagents and Anya Crane (Integrated Research Facility at Fort Detrick/National Institute of Allergy and Infectious Diseases/National Institutes of Health, Fort Detrick, Frederick, MD, USA) for critically editing the manuscript.

This work was supported by the National Institutes of Health grants AI160779 to AM and AKLL, AI164690 to AM, AI25271 to AM, GM104135 to AKLL for the development of ELTA-MS, AI078985 to GPL, GM07215 to VT, and F31GM143918 to IRU; the National Science Foundation GRFP DGE-1747503 to MPL, an H.I. Romnes Faculty Fellow funded by the Wisconsin Alumni Research Foundation and provided to AM; and a Vilas Faculty Mid-Career Investigator Award to AM. AM was a Burroughs Wellcome Fund Investigator in the Pathogenesis of Infectious Disease. All experiments were approved by the University of Wisconsin-Madison Institutional Biosafety Committee (IBC). The NIAID grant (R01AI25271) for the studies conducted was reviewed by the University of Wisconsin-Madison Dual Use Research of Concern (DURC) Subcommittee in accordance with the United States Government September 2014 DURC Policy and determined to meet the criteria of DURC. The University of Wisconsin-Madison Institutional Contact for Dual Use Research reviewed this manuscript and confirmed that the studies described herein do not meet the criteria of DURC. We acknowledge support of the shared instrumentation grant S10OD021844 from the Center for Proteomics Discovery at the Johns Hopkins University School of Medicine. This work was supported in part through Battelle Memorial Institute’s former prime contract with the U.S. National Institute of Allergy and Infectious Diseases (NIAID) under Contract No. HHSN272200700016I and Laulima Government Solutions, LLC current prime contract with NIAID under Contract No. HHSN272201800013C. SY, YC and JHK performed this work as former employees of Battelle Memorial Institute and current employees of Tunnel Government Services (TGS), a subcontractor of Laulima Government Solutions, LLC under Contract No. HHSN272201800013C.

The views and conclusions contained in this document are those of the authors and should not be interpreted as necessarily representing the official policies, either expressed or implied, of the U.S. Departments of the Army, Department of Defense, and Department of Health and Human Services, or of the institutions and companies affiliated with the authors, nor does mention of trade names, commercial products, or organizations imply endorsement by the U.S. Government.

## MATERIALS AND METHODS

### Cell lines

Authenticated stocks of the following cell lines were purchased from the American Type Culture Collection: human embryonic kidney 293T cells (CRL-3216), human lung A549 cells (CCL-185), Madin-Darby canine kidney cells (MDCK; CCL-34), chicken UMNSAH/DF-1 cells (CRL-3586), pig kidney PK(15) cells (CCL-33), and Brazilian free-tailed bat lung Tb 1 Lu cells (CCL-88). All cell lines were grown at 37°C with 5% CO_2_ in Dulbecco’s modified Eagle’s medium (DMEM) supplemented with 10% heat-inactivated fetal bovine serum. Cells were regularly tested and verified as free of *Mycoplasma* contamination using MycoAlert (Lonza LT07-318).

A549 PARP1 knockout cells were generated by co-transfecting the Cas9-expressing plasmid pMJ920 (Addgene 42234) and two different sgRNA-expressing plasmids targeting CCACCTCAACGTCAGGGTGC and TGGGTTCTCTGAGCTTCGGT creating a ∼35 bp deletion. Cells were transfected with TransIT-X2 (Mirus MIR6005) in 6-well plates using the recommended protocols. Twenty-four hours after transfection, clonal cell lines were generated by seeding on average 0.5 cells per well in a 96-well plate, expanding clones, and confirming the loss of PARP1 by blotting.

### Antibodies and ADPr detection reagents

Immunoblotting was performed with mouse α-tubulin DM1A (Sigma T9026), goat α-RNP (BEI Resources NR-3133), α-β actin (Proteintech 66009-1-Ig), mouse α-GFP B-2 (Santa Cruz Biotechnology SC-9996), and α-PARP1 (Cell Signaling 9542S). Immunoprecipitations were performed with mouse α-FLAG M2 (Sigma F1804) or α-NP H16-L10 (Bio X Cell BE0159) that was then captured on protein A agarose beads. ADP-ribosylation was detected by blotting with reagents specific for MAR, PAN or PAR (EMD Millipore MABE1076, MABE1016, and MABE1031, respectively)^53^. ADP-ribosylated proteins were enriched with GST-Af1521 that was expressed and purified from bacteria as described^35^. Where indicated, membranes were incubated overnight at room temperature with 1 M hydroxylamine (HAM; NH_2_OH) diluted in Tris-buffered saline and 0.5% Tween 20 before blotting for PAR.

### Transient and stable expression of PARPs

Vesicular stomatitis Indiana virus (VSIV) glycoprotein G-pseudotyped lentivirus was generated by transfecting 293T cells with the plasmids psPAX2 and pMD2.G, along with pLX304-PARP8 for constitutive PARP8 expression. A549 cells were transduced, selected with blasticidin (6 μg/ml), and a clonal population was isolated to create the PARP8-A549 cell line. A549 PARP1 knockout cells were complemented using the Sleeping Beauty transposon system^71^. Briefly, A549 PARP1 knockout cells were co-transfected with 0.2 μg of pSB100 and 1.8 μg of pSBbiBP-PARP1-V5 or PARP1 mutant plasmids. Cells were transfected with TransIT-X2 (Mirus MIR6005) in 6-well plate using the recommended protocols. Twenty-four hours after transfection, cells were selected with to 1μg/ml puromycin for 7 d. Expression of PARP1 or mutants was confirmed by blotting with anti-PARP1 or anti-V5-tag antibody. Transient PARP expression was achieved by transfecting 293T cells with plasmids expressing GFP-tagged PARP proteins^72^.

### Viruses and infections

Influenza A virus A/WSN/33 (H1N1; WSN), WSN encoding FLAG-tagged PB2 polymerase subunit (WSN-PB2-FLAG), and A/Puerto Rico/8/1934 (H1N1; PR8) were generated using the influenza A virus reverse genetics system^73,74^. Recombinant influenza reporter viruses WSN PA-Swap-2A-NanoLuc (WSN PASTN), WSN with the replication complex from A/green-winged teal/OH/175/1986 (H2N1) encoding PB2 S590/R591/K627 PASTN (S009 SRK PASTN), A/California/04/2009 PASTN (H1N1) (CA04 PASTN), and influenza B virus B/Brisbane/60/2008 (B/Brisbane PASTN) were generated as previously described^30,75,76^. Bat cell-adapted WSN was passaged 10 times in Tb 1 Lu cells as described in (26)^77^. PR8 that lacks NS1 (PR8Δ*NS1*) or encodes the RNA-binding mutant (NS1 K38A/R41A), as well as WSN that does not encode NS1 (WSNΔ*NS1*), were propagated and titered on MDCK cells overexpressing NS1-GFP.

293T cells were infected in virus growth medium (VGM; DMEM supplemented with penicillin/streptomycin, 25 mM HEPES, 0.3% BSA). A549 cells were infected in VGM with 0.25 μg/ml TPCK-trypsin or Opti-MEM I medium with 2% FBS and 0.25 μg/ml TPCK-trypsin. For detection of global ADP-ribosylation, A549 cells were infected at MOI 0.1, UMNSAH/DF-1 cells were infected in OptiMEM + 0.2% heat-inactivated FBS at MOI 1, and PK(15) and Tb 1 Lu cells were infected in OptiMEM + 2% FBS at MOI 1.

Viral gene expression was measured as described by inoculating cells with PASTN viruses diluted in VGM for 293T cells or in VGM with 0.25 μg/ml TPCK-trypsin for A549 cells^75,78^. Luciferase activity was measured 8 hpi using a Nano-Glo luciferase assay kit (Promega N1120). Multicycle replication infections were performed by inoculating cells with WSN PASTN at MOI 0.01 in VGM with 0.25 μg/ml TPCK-trypsin. Supernatants were collected at indicated times and titered by a Nano-Glo viral titer assay by inoculating MDCK cells with collected supernatants and measuring luciferase activity^75,78^.

### Detection of ADP-ribosylation

Cells were mocked-treated or infected with the indicated viruses. In some experiments, the PARG inhibitor PDD 00017273 (MedChemExpress HY-108360) was added to a final concentration of 3 μM for 24 hr before lysis. As a positive control, uninfected cells were exposed to 1 mM H_2_O_2_ for 10 min immediately before lysis. Cells were lysed in cold MOPS-RIPA buffer (16 mM MOPS pH 7.5, 150 mM NaCl, 1% deoxycholic acid, 0.2% SDS, 2% NP-40 with protease inhibitors, 1 μM ADP-HPD [Sigma 118415], and 40 μM PJ-34 [MilliporeSigma 528150]) or Tris-RIPA buffer (50 mM Tris pH 7.5, 150 mM NaCl, 0.5% deoxycholic acid, 0.1% SDS, 1% NP-40 with protease inhibitors and 1 μM ADP-HPD). The generic PARP inhibitor PJ-34 was included to prevent spurious ADP-ribosylation that can occur in lysates^79^. ADP-ribosylation was detected by blotting with purified MAR, PAN or PAR detection reagents^53^.

ADP-ribosylation was measured in the presence of the PARP inhibitors Olaparib (5 μM; MedChemExpress HY-10162), rucaparib (5 μM; MedChemExpress HY-10617A), AG14361 (10 μM; Sigma SML3081), XAV939 (5 μM; TOCRIS 3748), or PJ34 (4 μM; MedChemExpress HY-13688A). Stocks were prepared by dissolving inhibitors in DMSO. Cells were treated 1 h after infection.

### RNA sequencing

RNA-sequencing was previously reported^80^. Briefly, A549 cells were mock-infected or inoculated with WSN at an MOI of 0.02 in VGM with 0.25 μg/ml TPCK-trypsin. Infections were allowed to proceed for 24 h before RNA was harvested. In a separate condition, cells were mock-infected for 16 h followed by 8 h of interferon β (IFNβ) treatment (250 U/ml). Each condition was repeated in biological triplicate. RNA was extracted from cells using TRIzol (Invitrogen 15596026) per manufacturer’s specifications. RNA samples were submitted to Novogene and prepared for mRNA sequencing using NEBNext Ultra RNA Library Prep Kit for Illumina (New England Biolabs E7530). Reads were trimmed and mapped to a concatenated hg19-WSN genome using Spliced Transcripts Alignment to a Reference (STAR)^81^. Differential expression was evaluated with DESeq2^82^.

### Polymerase activity assay

293T cells were transfected with plasmids encoding WSN PA, PB1, PB2, and NP, a viral RNA-like firefly luciferase reporter, and a *Renilla* luciferase internal control reporter using TransIT-2020 (Mirus MIR 5400). 18 sites in NP above or near our 0.9 localization cutoff were selected for mutagenesis. Plasmids encoding mutant versions of viral proteins were created by PCR-mediated mutagenesis and verified by sequencing. In some experiments, additional proteins were co-expressed as indicated. Firefly and *Renilla* luciferase activities were assayed ∼24 h post-transfection. Firefly luciferase was normalized to *Renilla* luciferase within each sample.

### Sample preparation for proteomic analyses

Three distinct ELTA experiments were performed. In each experiment, each condition was performed in biological duplicate and samples were analyzed in technical triplicate. A549 cells were infected with WSN or PR8 at an MOI of 1 for 8 h and then treated with 1 mM H_2_O_2_ for 10 m or mock treated. In experiments with ΔNS1 viruses, infections with WSNΔNS1 or PR8ΔNS1 were allowed to proceed for 16 h such that viral protein levels were roughly equivalent with that in WT-infected cells. PDD was added to the cell culture prior to the end of the incubation. Cells were then placed on ice and quickly washed with cold PBS, which was then fully removed. Cells were lysed with freshly made guanidine-hydrochloride (GdnHCl) buffer heated to 99°C (6 M GdnHCl, 100 mM HEPES pH 8.0, 5 mM tris(2-carboxyethyl)phosphine [TCEP], 10 mM 2-chloroacetamide). Cells were scraped from the dish and transferred to an Eppendorf tube where they were incubated for an additional 10 min at 99°C. Samples from the same biological replicate were pooled, flash frozen in liquid nitrogen, and stored at -80°C until processing. Samples were thawed and total protein concentration of lysates was determined by Bradford assay. Protein digestion reactions were prepared by diluting lysates containing 10 mg total protein 6-fold in 100 mM Tris-HCl pH 8.0 and adding 100 µg Trypsin (Thermo Fisher 90058) and 100 µg LysC (Wako 129-02543). Samples were incubated overnight at 37°C while shaking at 200 rpm. Next, 1 M triethylammonium acetic acid (TEAA) pH 7.5 was added to peptide solutions to a final concentration of 100 mM before clarifying peptide solutions by centrifugation at 4,000 rpm for 30 min at 4°C. 1 mg SepPak t-C18 Classic cartridges (Waters) were conditioned with 80% acetonitrile (ACN) and equilibrated with 100 mM TEAA pH 7.5 before loading peptide solutions. Peptides were washed with 100 mM TEAA pH 7.5 and eluted with 40% ACN. Samples were dried down to completion by vacuum centrifugation.

### Enrichment of ADP-ribosylated peptides

ADP-ribosylated peptides were resuspended in MilliQ H_2_O and enriched using the ELTA-MS proteomics workflow^36,37^. First, peptides were incubated with 100 µM N6-(6-Azido)hexyl-dATP (Jena Bioscience CLK-NU-002), 20 µg/mL low molecular weight poly(I:C) (Invivogen tlrl-picw), and 20 µg/mL human 2’-5’-oligoadenylate synthase 1 (OAS1; prepared in-house as in ^36^) in buffer containing 100 mM Tris-HCl pH 7.5, 20 mM magnesium acetate, and 2.5 mM dithiothreitol (DTT) for 1 h at 37°C while shaking at 800 rpm. 100 µL of 50% dibenzocyclooctyne (DBCO)-agarose (Click Chemistry Tools 1034) equilibrated in 1x PBS was added to each OAS1 reaction before rotating end-over-end at room temperature for 1 h. The samples were centrifuged to pellet the agarose resin and the supernatant was discarded. The agarose resin was washed three times with 5 M NaCl, three times with 20% ACN, and three times with 1x PBS. After washing the resin twice in phosphodiesterase buffer containing 100 mM HEPES pH 8.0 and 15 mM MgCl_2_, the samples were transferred to new Eppendorf tubes. Human nudix 16 (NudT16) was prepared as described^83^, 5 μg was added to each sample, and the volume was brought up to 200 µL with phosphodiesterase buffer. Samples were incubated for 2h at 37°C while shaking at 1400 rpm. The samples were centrifuged to pellet the agarose resin and 150 µL of the eluate was transferred to new Eppendorf tubes. 10% trifluoroacetic acid (TFA) was added to samples to a final concentration of 1% TFA, and the samples were loaded onto homemade Stage-Tips that were conditioned with 80% ACN/0.1% TFA and equilibrated with 5% ACN/0.1% TFA. Peptide were washed with 0.1% TFA, eluted with 40% ACN/0.1% TFA, and dried down to completion by vacuum centrifugation.

### LC-MS/MS analysis of enriched peptides

All enriched peptide samples were analyzed on a Thermo Orbitrap Fusion Lumos mass spectrometer coupled to an EASY-nLC 1200 system (Thermo) with a column type and stationary phase. LC Buffer A was 0.1% formic acid (FA), LC Buffer B was 0.1% FA/95% ACN, and the column oven was maintained at 40°C. Peptides were separated with a 95-min gradient from 8% to 27% Buffer B, followed by an increase to 50% Buffer B over 5 min, another increase to 90% Buffer B over 5 min, and finally a 10-min washing block. MS data were collected with a spray voltage of 2.4 kV, capillary temperature of 200 °C, and RF lens of 30%. MS1 scans were performed at a resolution of 120 k, a scan range of 300 to 1800 m/z, a maximum injection time of 50 ms, and an automatic gain control (AGC) target of 1,000,000 charges. Precursor ions were isolated with a width of 1.6 m/z and an AGC target of 50,000 charges. Higher-energy collisional dissociation (HCD) fragmentation was performed with a normalized collision energy of 25% and a resolution of 30 k.

### MS data analysis

Thermo .RAW files were converted to .mzML files using MSConvert from the ProteoWizard library^84^. Database searching was performed in FragPipe (v 20.0) with MSFragger (v 3.8), IonQuant (v 1.9.8), and Philosopher (v 5.0.0)^85–87^. The UniProt FASTA file for PR8 proteins PA, PA-X, PB1, PB1-F2, PB2, NP, HA, NA, M1, M2, NS1, and NEP/NS2 were downloaded from influenza A virus (strain A/Puerto Rico/8/1934 H1N1; Taxon ID 211044). The UniProt FASTA file for WSN proteins PA, PB1, PB2, NP, HA, NA, M1, M2, and NS1 were downloaded from influenza A virus (A/WSN/1933 H1N1; Taxon ID 382835) and WSN proteins PA-X and NEP/NS2 from influenza A virus (strain A/Wilson-Smith/1933 H1N1; Taxon ID 381518). The UniProt FASTA file for the human proteome was downloaded from UniProt Proteome ID UP000005640. The four FASTA files were manually concatenated in Python, and then decoys and contaminant proteins were added in FragPipe. A closed search in nonlabile mode was performed using the default FragPipe parameters with exceptions specified below. Fully enzymatic digestion used Trypsin and LysC rules. Variable modifications included 212.0086 (phospho-ribosylation) on DESKTYRCH (max 2 per peptide), 15.9949 (oxidation) on M (max 3 per peptide), 42.0106 (acetylation) on protein N-terminus (max 1), and -0.98401 (amidation) on protein C-terminus (max 1). The fixed modification was indicated as 57.02146 (carbamidomethylation) on C. The activation type filter was HCD and fragment ion series included b and y. Peptide-spectrum match (PSM) validation was performed with Percolator, PTM site localization was performed with PTMProphet, and protein inference was performed via ProteinProphet^88–90^. “Match between runs” (MBR) was turned on and the MBR ion false discovery rate (FDR) was set to 1%. “Normalize intensity across runs” was turned on and “peptide-protein uniqueness” was set to unique and razor peptides.

### MS data filtering and visualization

All downstream analyses of the DESKTYRCH_212.0086.tsv and combined_ion.tsv output files of the FragPipe search and plotting of the processed data were performed with homemade scripts in the statistical environment R (v. 4.3.2). Incorrectly assigned phospho-ribose sites on C were removed from the DESKTYRCH_212.0086 dataframe by comparing it to the carbamidomethylation sites on C in the combined_ion dataframe. Site indices [protein ID_amino acid_residue] were calculated for the combined_ion data frame. Peptides with one phospho-ribose were labeled as “pep_one-mod” and peptides with two phospho-riboses were labeled as “pep_multi-mod” and then two separate index rows were made for each multiply-modified peptide. The MS data (RT, m/z, charge, PSMs, etc.) were copied to each separate index row associated with the multiply-modified peptide. To ensure reproducibility across technical replicates and filter for modified peptide identifications with direct MS evidence, we developed a formula to calculate a score for each site index. MS/MS identifications were assigned an arbitrary value of 3 and MBR identifications were assigned an arbitrary value of 1. For each biological replicate, we scored all cases in which the site was identified in at least two technical replicates. The score equaled the sum of MS/MS identifications multiplied by 3 plus the sum of MBR identifications multiplied by 1. Modified peptide identifications were filtered for scores greater than or equal to 4, indicating at least one technical replicate containing direct MS/MS evidence of the modified peptide and another replicate with either matching or direct evidence. For plotting, modified peptide identifications were condensed into unique site indices. For PSMs of site indices derived from multiple modified peptide sequences, the PSMs for each modified peptide associated with that site index were summed. For PSMs of modified proteins, the PSMs for every modified peptide/site index associated with that protein were summed, excluding the ones labeled “pep_multi-mod” to avoid double counting.

For the clustering heatmaps, unique site indices were filtered for localization probabilities greater than or equal to 0.9. Then, identifications with scores greater than or equal to 4 were assigned a value of one (present) and identifications that were unmatched or had a score of less than 4 were assigned a value of zero (absent). Clustering heatmaps were generated using the pheatmap function in R (v. 4.3.2) and the Jaccard (“binary”) clustering distance for categorical data. Heatmaps showing the number of sites, max score, or number of PSMs were generated using ggplot2 in R.

For Venn diagrams, modified site indices were filtered for localization probabilities greater than or equal to 0.9, whereas modified proteins were filtered less stringently for localization probabilities greater than or equal to 0.6 in order to increase the number of modified peptide identifications, since ADPr is a labile modification during HCD fragmentation and thus can be difficult to localize. Venn diagrams were generated using InteractiVenn^91^.

### Statistics

All experiments were repeated with at least three independent biological replicates, with at least three technical replicates for each experiment. Representative replicates are shown as the mean ± standard deviation. Pairwise comparisons were made using a two-tailed Student’s t-test. Multiple comparisons were performed with an ANOVA followed by an *ad hoc* Dunnett’s multiple comparisons test. GO enrichment was performed in gProfiler (2.0) and ShinyGO (0.8)^92,93^.

## RESOURCE AVAILABILITY

### Lead Contact Statement

Further information and requests for resources and reagents should be directed to and will be fulfilled by the Lead Contact, Andrew Mehle.

### Materials Availability

All unique reagents generated in this study are available from the Lead Contact with a completed Materials Transfer Agreement.

### Data availability

RNA-seq data are available as part of BioProject PRJNA667475. Mass spectra raw files are accessible on the MassIVE database under accession code MSV000095351 with a ProteomeXchange ID PXD053989. Other raw data are available in Figure S10 and Table S5.

