## Supplementary material for "Global remodeling of ADP-ribosylation by PARP1 suppresses influenza A virus infection": Sup Fig 1

### Supplemental Information titles and legends

**Figure S1. Activation of PARP gene expression following infection or interferon β treatment. Related to Figure 1**
(A) Expression level of each PARP (log2[FPKM]) in human lung A549 cells. High- and low-expressing genes are plotted separately for clarity.
(B) Fold induction of each PARP during infection with WSN or IFNβ treatment of A549 cells was calculated relative to mock-infected A549 cells. PARP15 was excluded because it does not express in mock-infected A549 cells.

For all, data are mean of n=3 ± SD. Treatment conditions were compared to control for each PARP using a one-way ANOVA with post-hoc Tukey HSD test. *$p<0.05$, **$p<0.01$.

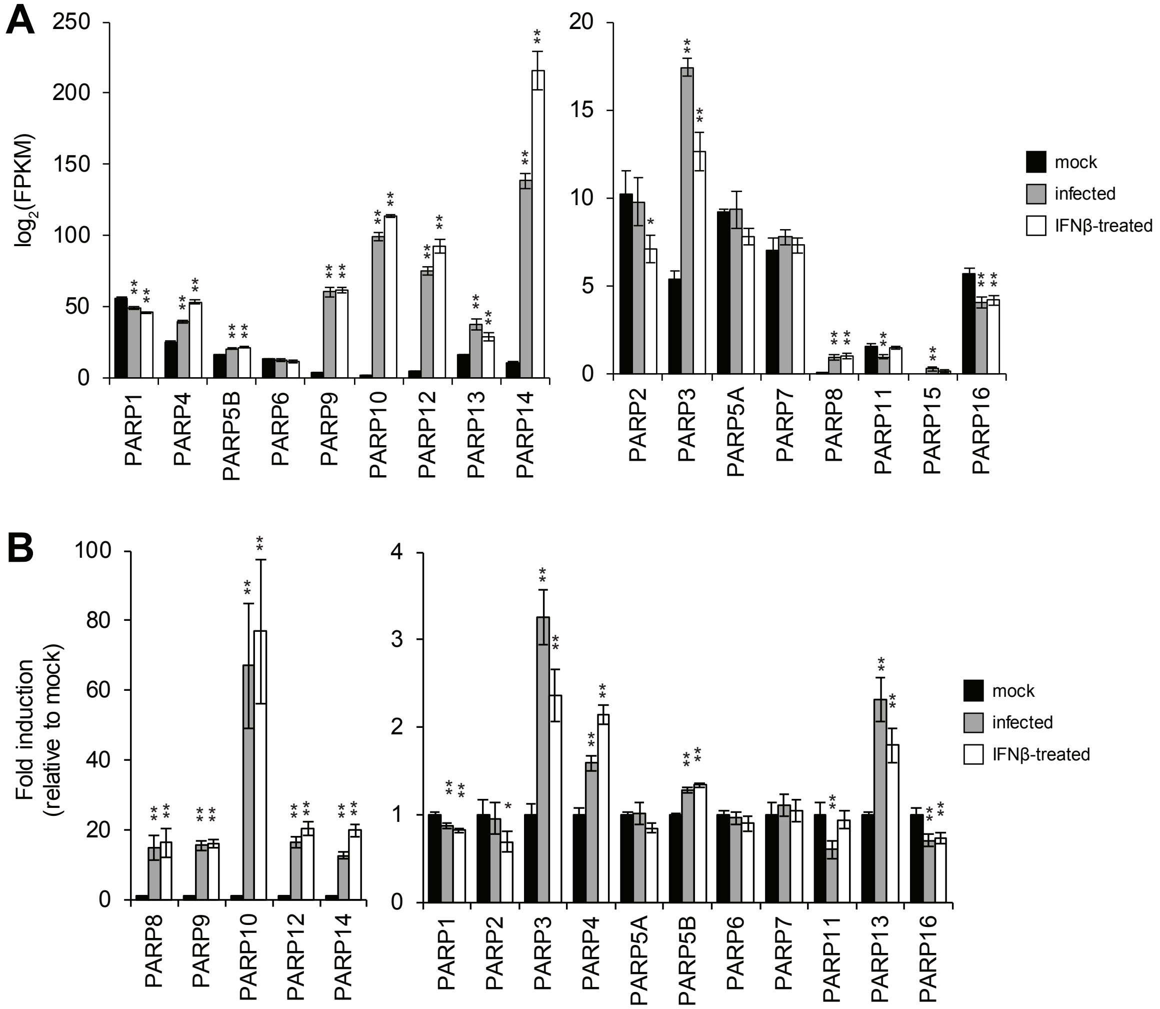
A
log2(FPKM)
250
200
150
100
50
0
PARP1
PARP4
PARP5B
PARP6
PARP9
PARP10
PARP12
PARP13
PARP14
20
15
10
5
0
PARP2
PARP3
PARP5A
PARP7
PARP8
PARP11
PARP15
PARP16
mock
infected
IFNβ-treated
B
Fold induction
(relative to mock)
100
80
60
40
20
0
PARP8
PARP9
PARP10
PARP12
PARP14
4
3
2
1
0
PARP1
PARP2
PARP3
PARP4
PARP5A
PARP5B
PARP6
PARP7
PARP11
PARP13
PARP16
mock
infected
IFNβ-treated
