## Supplementary material for "Global remodeling of ADP-ribosylation by PARP1 suppresses influenza A virus infection": Sup Fig 2

**Figure S2. PARP expression interferes with viral infection and polymerase activity. Related to Figure 1**

(A) Multicycle replication of influenza A virus WSN in human lung A549 cells and those stably expressing PARP8 or IFITM3.

(B) PARP8 expression suppresses viral gene expression. A549 cells expressing PARP8, IFITM1, IFITM2, or IFITM3 were infected with WSN PASTN and viral gene expression assayed.

(C) Viral gene expression for the indicated strains in A549 cells or cells stably expressing PARP8.

For all, data are presented as mean of 3 to 4 replicates ± SD. Multiple comparisons were made in A and B using a one-way ANOVA with post-hoc Dunnett's test when compared to the control. Pairwise comparisons in C were performed with a Student's two-tailed t-test. ** = $p<0.01$, **** = $p<0.0001$.

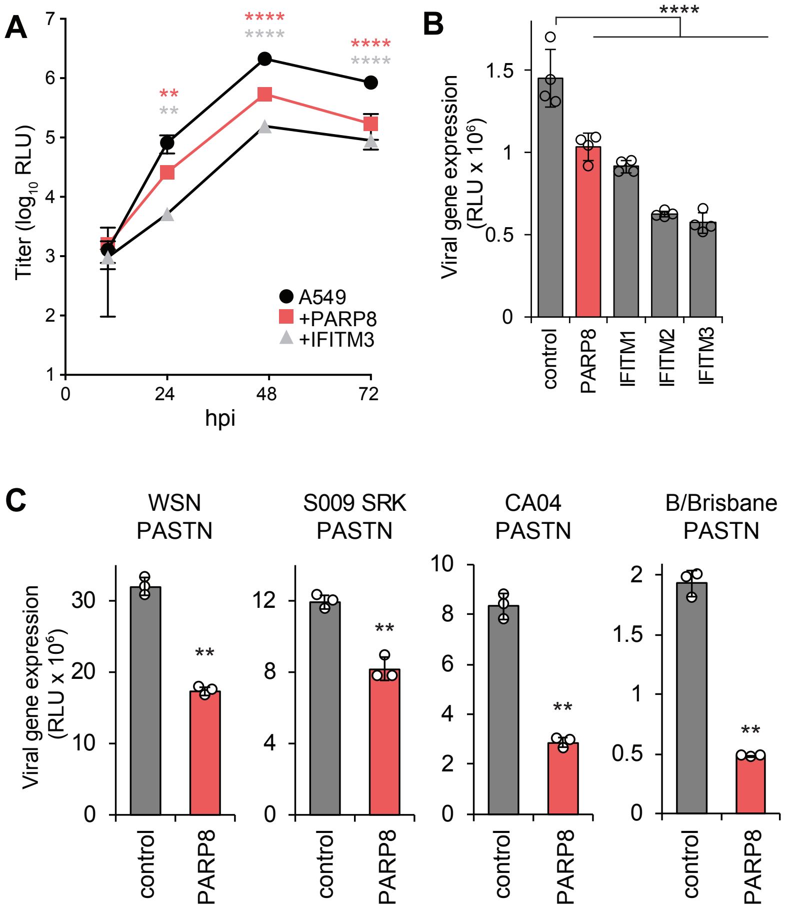
A
Titer (log10 RLU)
hpi
A549
+PARP8
+IFITM3
B
Viral gene expression (RLU x 10^6)
control
PARP8
IFITM1
IFITM2
IFITM3
C
WSN PASTN
S009 SRK PASTN
CA04 PASTN
B/Brisbane PASTN
Viral gene expression (RLU x 10^6)
control
PARP8
