## Supplementary material for "Global remodeling of ADP-ribosylation by PARP1 suppresses influenza A virus infection": Sup Fig 3

**Figure S3. Influenza A virus NP is ADP-ribosylated. Related for Figures 1-2**
(A) Human lung A549 cells were infected with WSN. ADP-ribosylated proteins were affinity purified with recombinant Af1521 or the binding mutant G42E. Influenza A virus NP was detected in the input and purified fractions by blotting.
(B) ADP-ribosylation sites identified in Figure 2 were mapped onto domains in NP. Domains and RNA-binding residues were defined based on prior structural analyses[55–57]. The figure is fully zoomable with modified sites aligned with the NP (WSN) amino acid sequence shown above the cartoon. NLS, nuclear localization signal.
(C) ADP-ribosylation sites identified in Figure 2 are shown on the structure of an NP monomer from WSN (PDB 2IQH). The surface of NP is colored by electrostatic potential, except for ADP-ribosylation sites that are colored yellow. ADP-ribosylation sites whose mutation increases polymerase activity in Figure 2F are highlighted. S69, T92, and S367 contact RNAs that are bound by NP[55]. E294 is part of the inter-strand interface[94].

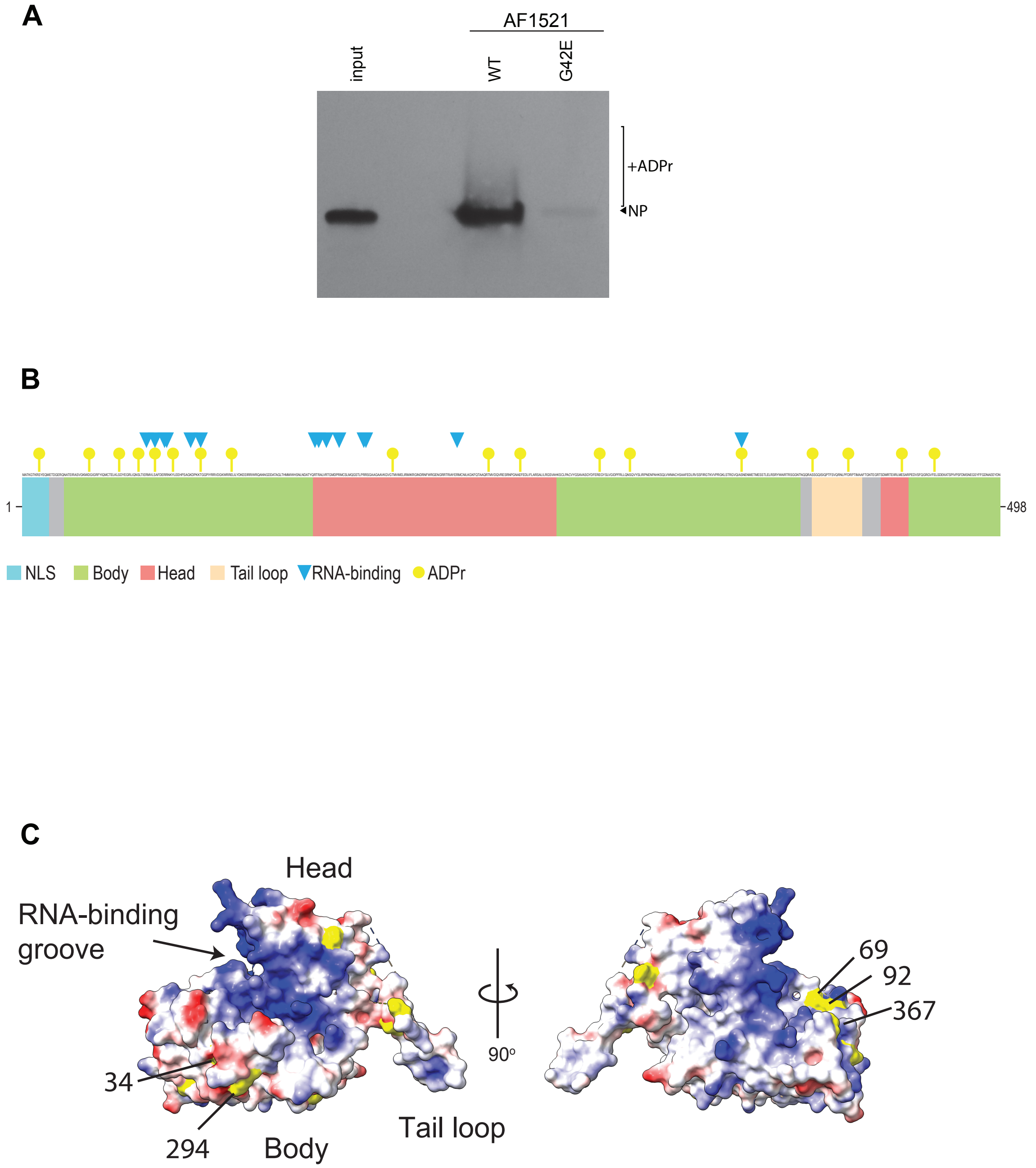
A
AF1521
input
WT
G42E
+ADPr
NP
B
1
498
NLS
Body
Head
Tail loop
RNA-binding
ADPr
C
Head
RNA-binding groove
34
294
Body
Tail loop
90°
69
92
367
