## Supplementary material for "Global remodeling of ADP-ribosylation by PARP1 suppresses influenza A virus infection": Sup Fig 4

**Figure S4. Reproducibility of ELTA. Related to Figure 2**

(A) Venn diagram of ADP-ribosylated proteins and sites as in Figure 2B. ELTA was performed on two independent biological replicates per condition, each analyzed in technical triplicate. Cells were infected with WSN, treated with $H_2O_2$, or mock-treated. Overlap of technical (left) and biological replicates (right) are shown at both the protein and peptide level.

(B) Enrichment of GO biological processes in the core ADP-ribosylome shared by all conditions.

(C) Comparison of ADP-ribosylome between A549 and previously reported $H_2O_2$-treated HeLa cells[37].

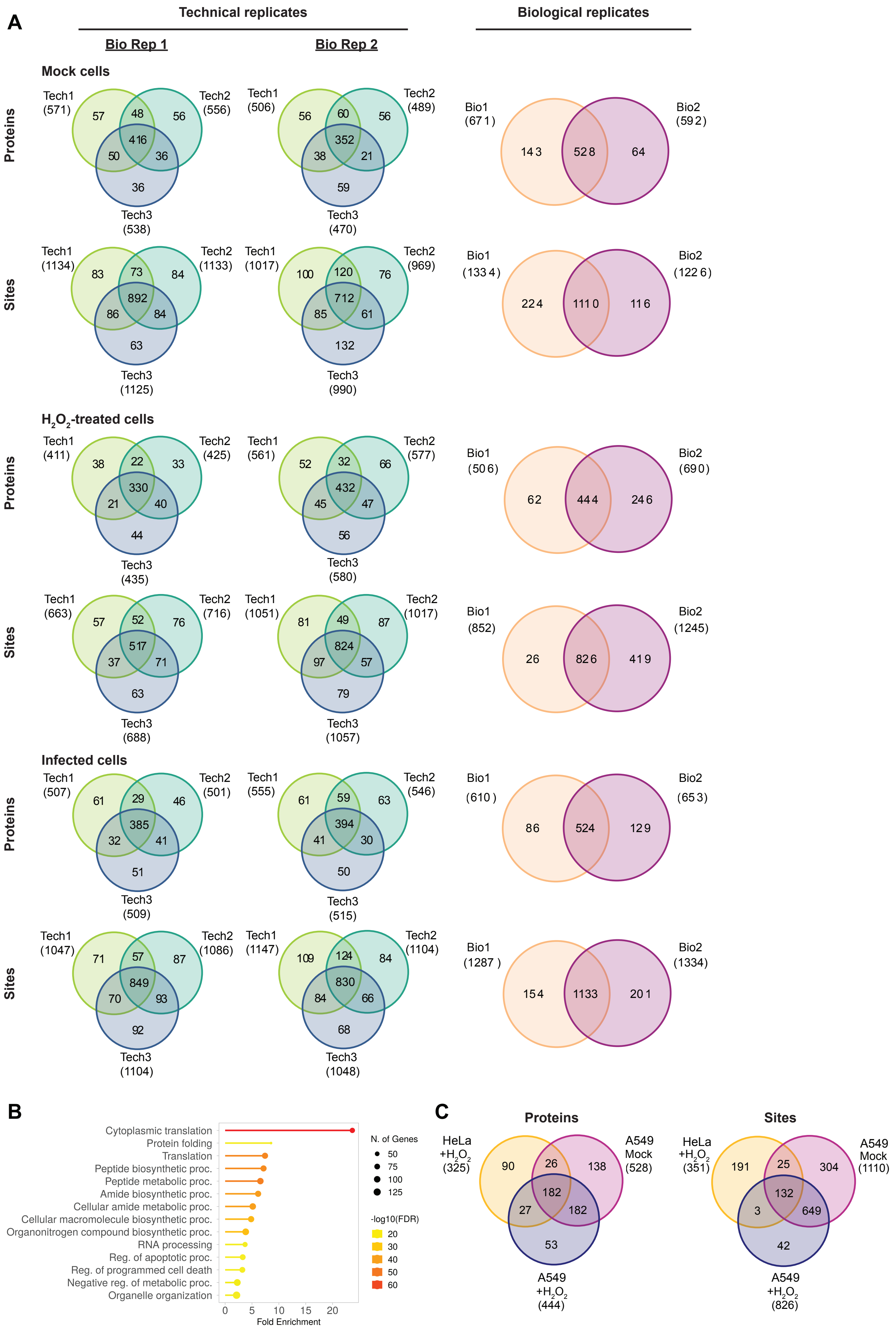
A
Technical replicates
Biological replicates
Bio Rep 1
Bio Rep 2
Mock cells
Proteins
Tech1 (571)
Tech2 (556)
Tech3 (538)
57
48
56
416
50
36
36
Tech1 (506)
Tech2 (489)
Tech3 (470)
56
60
56
352
38
21
59
Bio1 (671)
Bio2 (592)
143
528
64
Sites
Tech1 (1134)
Tech2 (1133)
Tech3 (1125)
83
73
84
892
86
84
63
Tech1 (1017)
Tech2 (969)
Tech3 (990)
100
120
76
712
85
61
132
Bio1 (1334)
Bio2 (1226)
224
1110
116
$H_2O_2$-treated cells
Proteins
Tech1 (411)
Tech2 (425)
Tech3 (435)
38
22
33
330
21
40
44
Tech1 (561)
Tech2 (577)
Tech3 (580)
52
32
66
432
45
47
56
Bio1 (506)
Bio2 (690)
62
444
246
Sites
Tech1 (663)
Tech2 (716)
Tech3 (688)
57
52
76
517
37
71
63
Tech1 (1051)
Tech2 (1017)
Tech3 (1057)
81
49
87
824
97
57
79
Bio1 (852)
Bio2 (1245)
26
826
419
Infected cells
Proteins
Tech1 (507)
Tech2 (501)
Tech3 (509)
61
29
46
385
32
41
51
Tech1 (555)
Tech2 (546)
Tech3 (515)
61
59
63
394
41
30
50
Bio1 (610)
Bio2 (653)
86
524
129
Sites
Tech1 (1047)
Tech2 (1086)
Tech3 (1104)
71
57
87
849
70
93
92
Tech1 (1147)
Tech2 (1104)
Tech3 (1048)
109
124
84
830
84
66
68
Bio1 (1287)
Bio2 (1334)
154
1133
201
B
Cytoplasmic translation
Protein folding
Translation
Peptide biosynthetic proc.
Peptide metabolic proc.
Amide biosynthetic proc.
Cellular amide metabolic proc.
Cellular macromolecule biosynthetic proc.
Organonitrogen compound biosynthetic proc.
RNA processing
Reg. of apoptotic proc.
Reg. of programmed cell death
Negative reg. of metabolic proc.
Organelle organization
0 5 10 15 20
Fold Enrichment
N. of Genes
50
75
100
125
-log10(FDR)
20
30
40
50
60
C
Proteins
HeLa $+H_2O_2$ (325)
A549 Mock (528)
A549 $+H_2O_2$ (444)
90
26
138
182
27
182
53
Sites
HeLa $+H_2O_2$ (351)
A549 Mock (1110)
A549 $+H_2O_2$ (826)
191
25
304
132
3
649
42
