## Supplementary material for "Global remodeling of ADP-ribosylation by PARP1 suppresses influenza A virus infection": Sup Fig 5

**Figure S5. ADP-ribosylation in the absence of NS1**
A549 cells were infected with PR8, PR8Δ*NS1*, mock-infected, or treated with $H_2O_2$. ADP-ribosylation sites were identified by ELTA-MS.
(A) Heat map clustering unique ADP-ribosylation sites of human and viral proteins present in each biological replicate.
(B) Venn diagram of modified proteins (left) and sites (right) identified in (A)
(C) Heatmap of the number of ADP-ribosylation sites identified on viral proteins during infection.
(D) Distribution of modifications detected on the specified amino acids for each condition and replicate.

For each condition, ELTA-MS was performed in technical triplicate on two independent biological replicates.

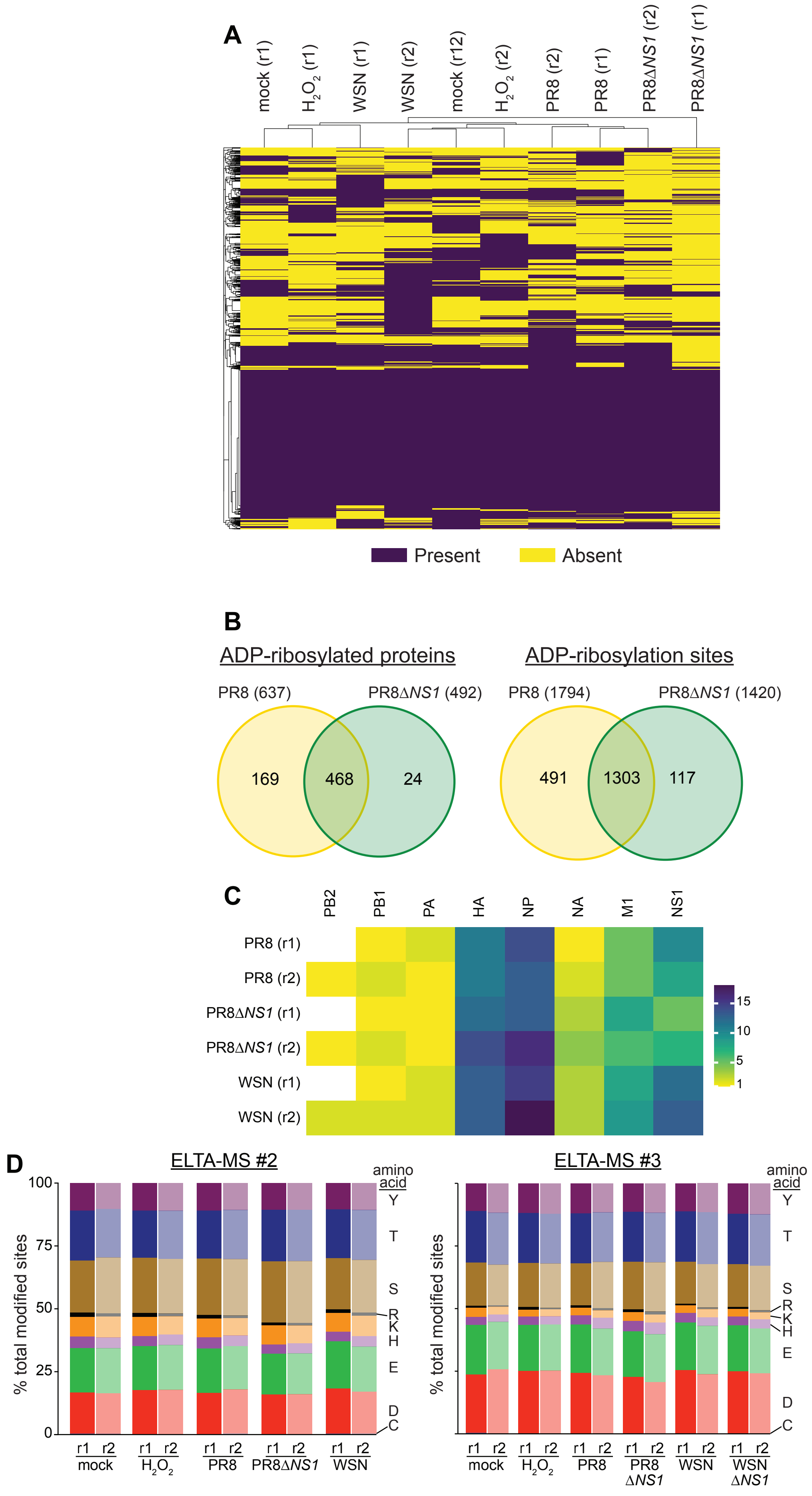
A
mock (r1)
H2O2 (r1)
WSN (r1)
WSN (r2)
mock (r12)
H2O2 (r2)
PR8 (r2)
PR8 (r1)
PR8ΔNS1 (r2)
PR8ΔNS1 (r1)
Present
Absent
B
ADP-ribosylated proteins
PR8 (637)
PR8ΔNS1 (492)
169
468
24
ADP-ribosylation sites
PR8 (1794)
PR8ΔNS1 (1420)
491
1303
117
C
PB2
PB1
PA
HA
NP
NA
M1
NS1
PR8 (r1)
PR8 (r2)
PR8ΔNS1 (r1)
PR8ΔNS1 (r2)
WSN (r1)
WSN (r2)
15
10
5
1
D
ELTA-MS #2
% total modified sites
100
75
50
25
0
amino acid
Y
T
S
R
K
H
E
D
C
r1 r2
mock
H2O2
PR8
PR8ΔNS1
WSN
ELTA-MS #3
% total modified sites
amino acid
Y
T
S
R
K
H
E
D
C
r1 r2
mock
H2O2
PR8
PR8 ΔNS1
WSN
WSN ΔNS1
