## Supplementary material for "Global remodeling of ADP-ribosylation by PARP1 suppresses influenza A virus infection": Sup Fig 6

**Figure S6. Reproducibility of ELTA (related to Figures 2 and 5)**
(A) Venn diagram of ADP-ribosylated proteins and sites for each infection condition.
(B) Venn diagram showing a shared core of ADP-riboyslated proteins in infected cells. Data are obtained cells infected with WSN or PR8 in ELTA-MS datasets #2 and #3.

ELTA was performed on two independent biological replicates per condition, each analyzed in technical triplicate. Cells were infected, treated with $H_2O_2$, or mock-treated.

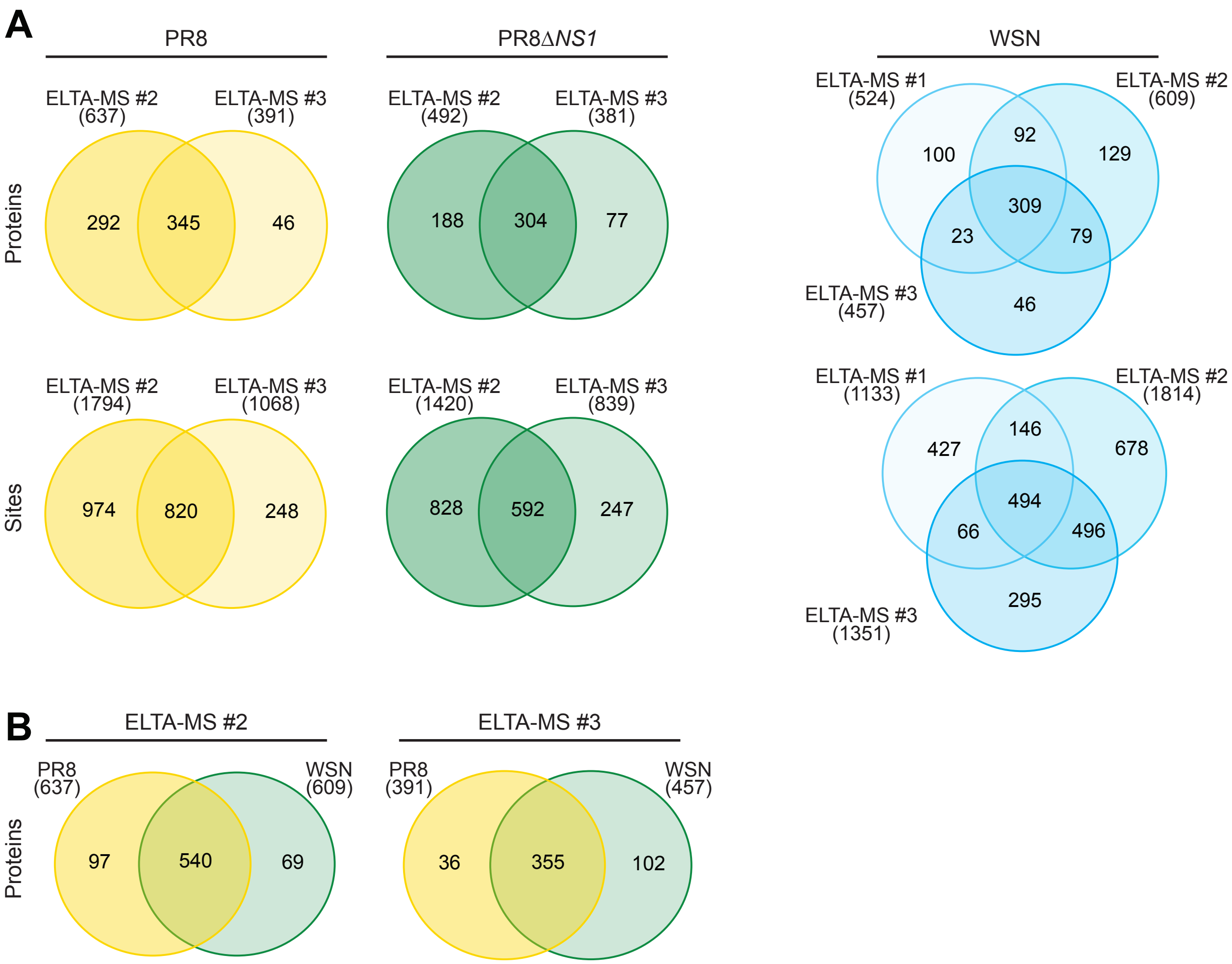
A
PR8
PR8Δ*NS1*
WSN
Proteins
Sites
ELTA-MS #2
(637)
ELTA-MS #3
(391)
292
345
46
ELTA-MS #2
(492)
ELTA-MS #3
(381)
188
304
77
ELTA-MS #1
(524)
ELTA-MS #2
(609)
ELTA-MS #3
(457)
100
92
129
309
23
79
46
ELTA-MS #2
(1794)
ELTA-MS #3
(1068)
974
820
248
ELTA-MS #2
(1420)
ELTA-MS #3
(839)
828
592
247
ELTA-MS #1
(1133)
ELTA-MS #2
(1814)
ELTA-MS #3
(1351)
427
146
678
494
66
496
295
B
ELTA-MS #2
ELTA-MS #3
Proteins
PR8
(637)
WSN
(609)
97
540
69
PR8
(391)
WSN
(457)
36
355
102
