## Supplementary material for "Global remodeling of ADP-ribosylation by PARP1 suppresses influenza A virus infection": Sup Fig 7

**Figure S7. PARP1 mediates PARylation during infection. Related to Figure 7**
(A) PARylating activity of PARP1 is required for infection-induced ADP-ribosylation. Human lung A549 *PARP1* KO cells were complemented with PARP1-V5 or the PARylating mutants E988A or E988K. Cells were infected with WSNΔNS1 or mock treated and whole cell extracts were used for blotting. Lysates were probed for PARP1, V5, NP as a marker of infection, and β-actin as a loading control.
(B) Caspase-cleavage of PARP1 is important for PARylation during infection. Human lung A549 *PARP1* KO cells were complemented with PARP1-V5 or PARP1 D214A with a mutated caspase-cleavage site. Cells were infected with PR8Δ*NS1* or mock treated and whole cell extracts were used for blotting. Lysates were probed for PARP1, V5, NP as a marker of infection, and β-actin as a loading control.

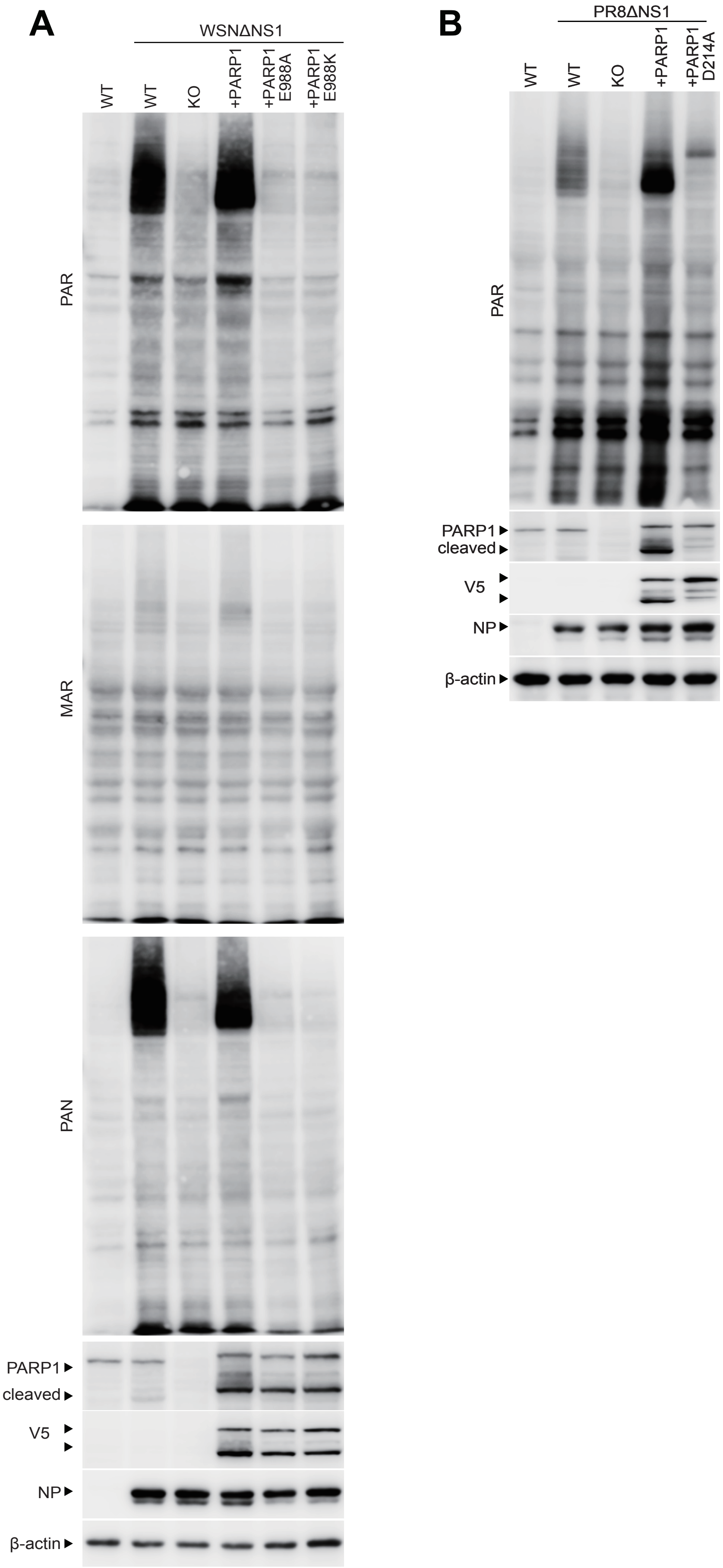
A
WSNΔNS1
WT
WT
KO
+PARP1
+PARP1 E988A
+PARP1 E988K
PAR
MAR
PAN
PARP1
cleaved
V5
NP
β-actin
B
PR8ΔNS1
WT
WT
KO
+PARP1
+PARP1 D214A
PAR
PARP1
cleaved
V5
NP
β-actin
