## Supplementary figures and images for "Global remodeling of ADP-ribosylation by PARP1 suppresses influenza A virus infection"

### Sup Fig 8

**Figure S8 Diagram of ADPr sites identified on viral proteins from A/WSN/33**

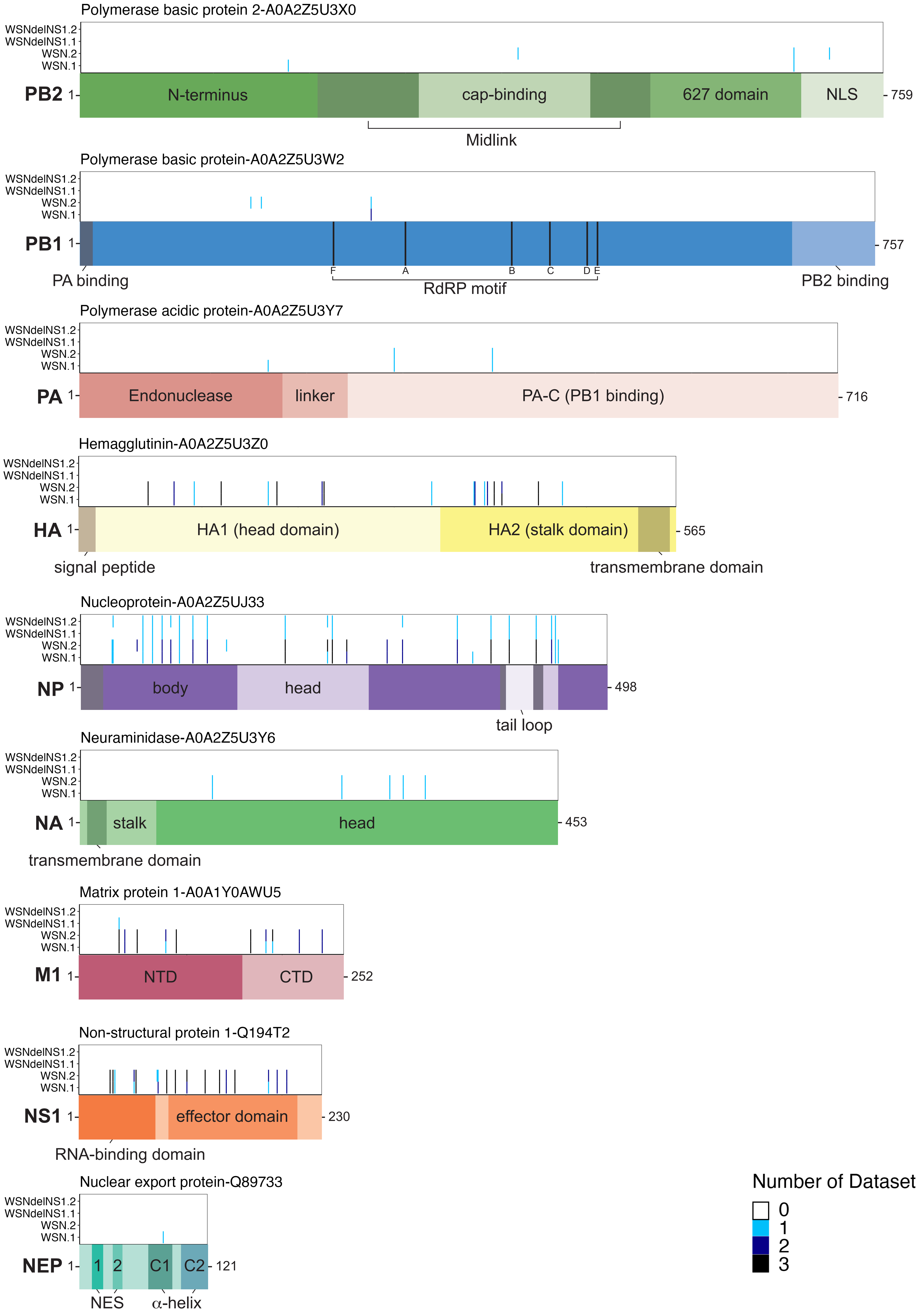

### Sup Fig 9

**Figure S9 Diagram of ADPr sites identified on viral proteins from A/PR8/34**

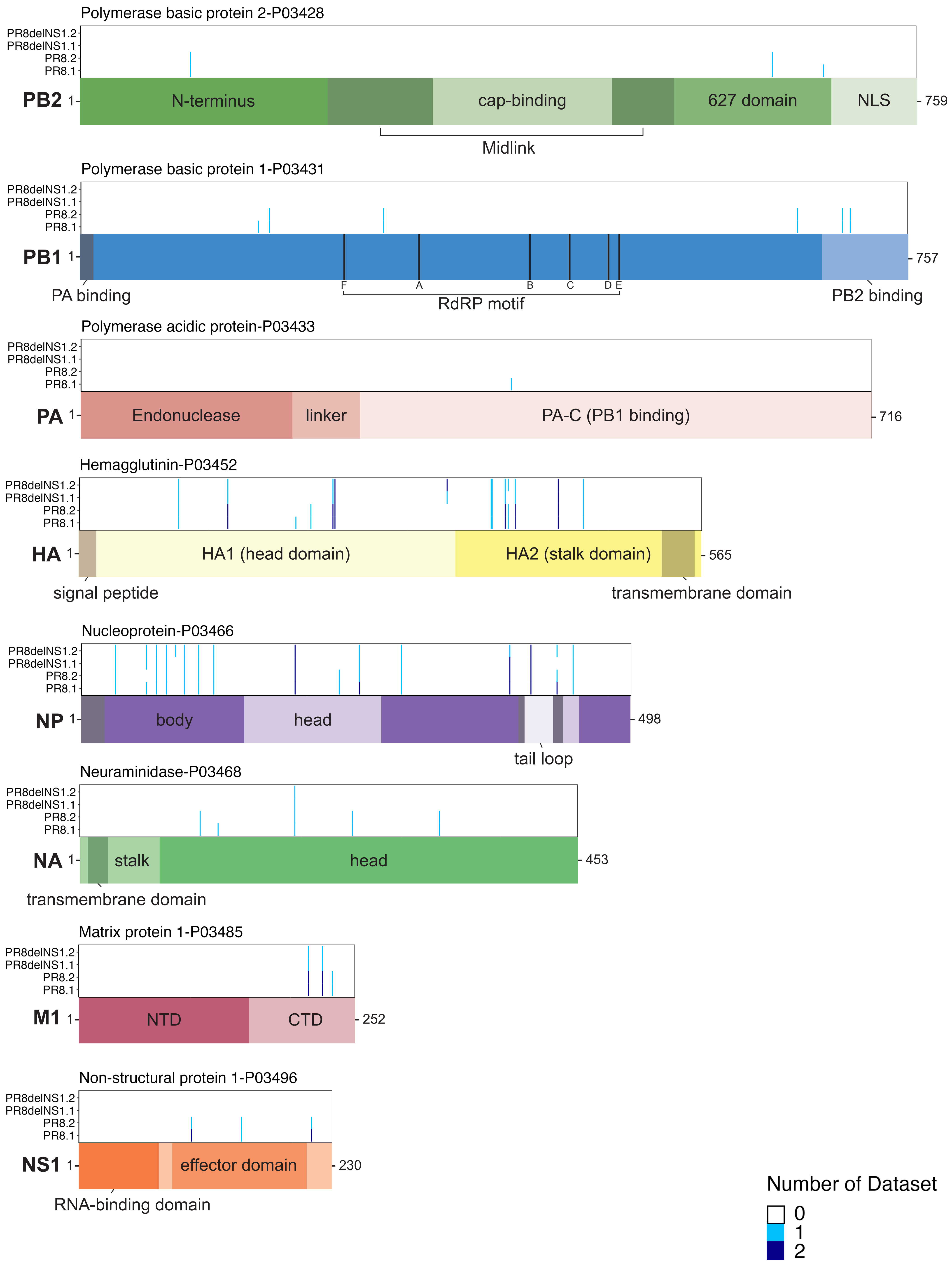
