## Supplementary material for "Global remodeling of ADP-ribosylation by PARP1 suppresses influenza A virus infection": Sup Fig 10

#### Figure S10 Uncropped blots

Figure 1. 1D

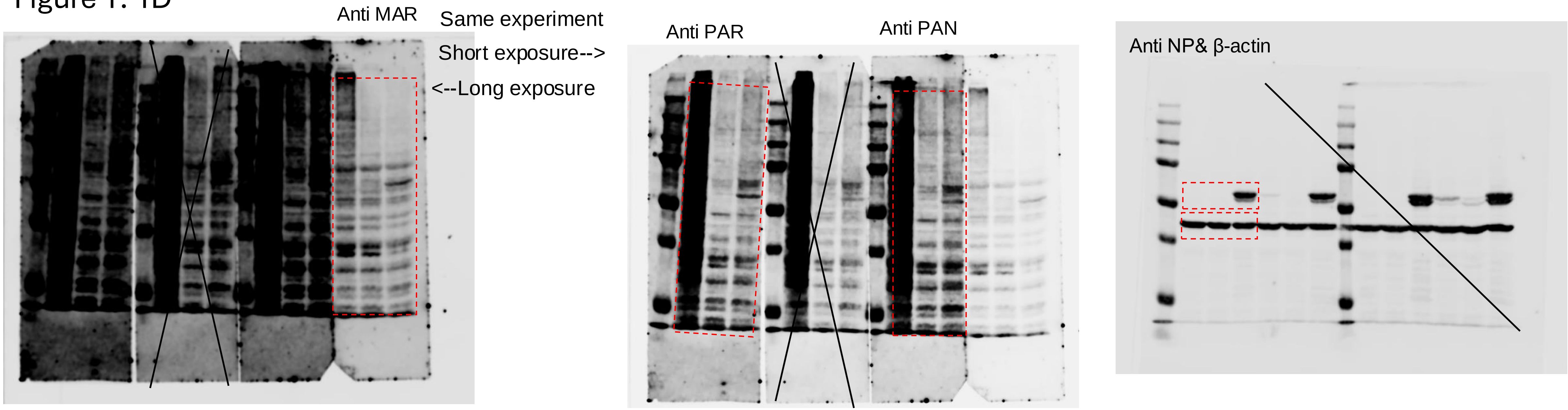


1E left

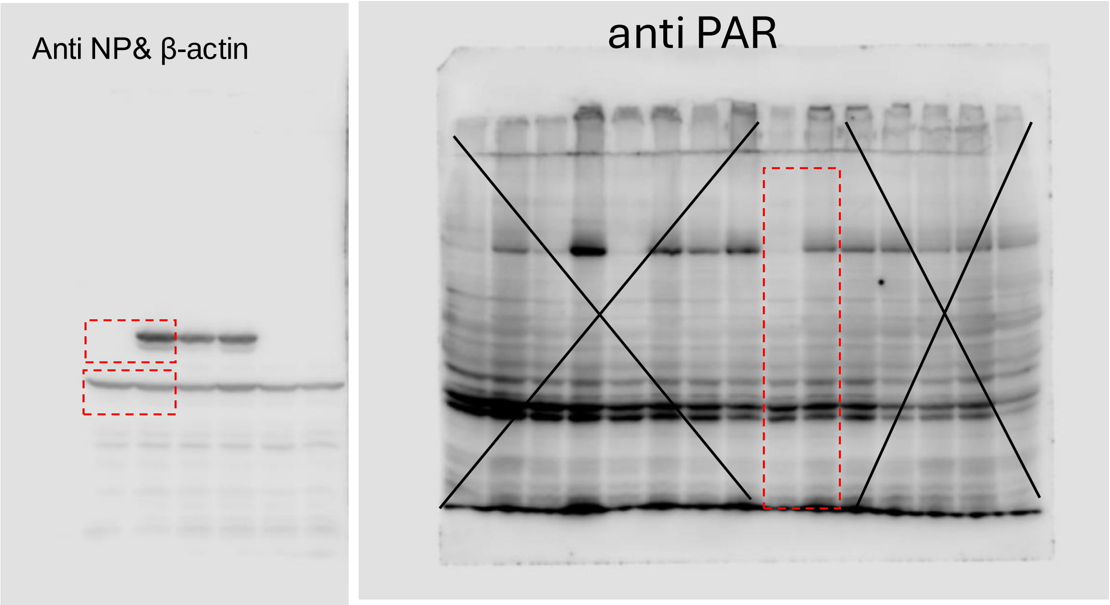


1E right

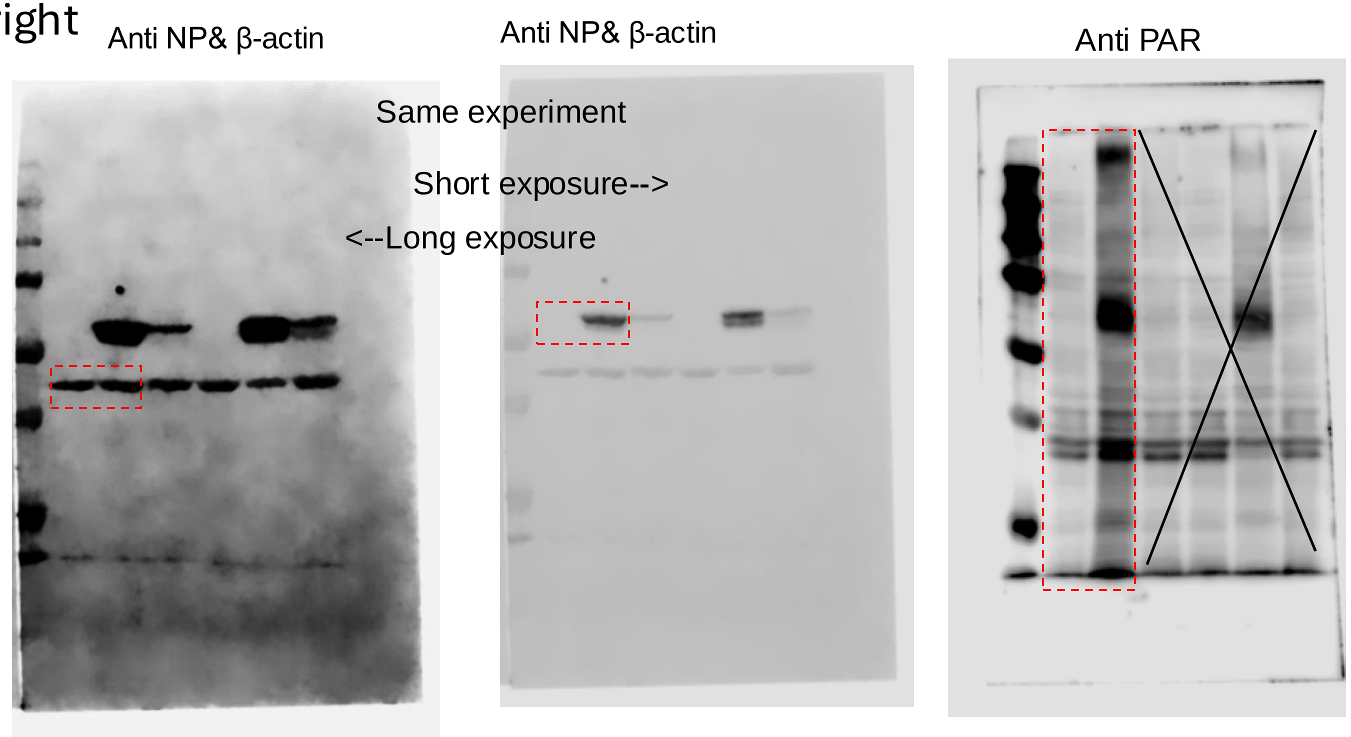

Figure 1. 1F

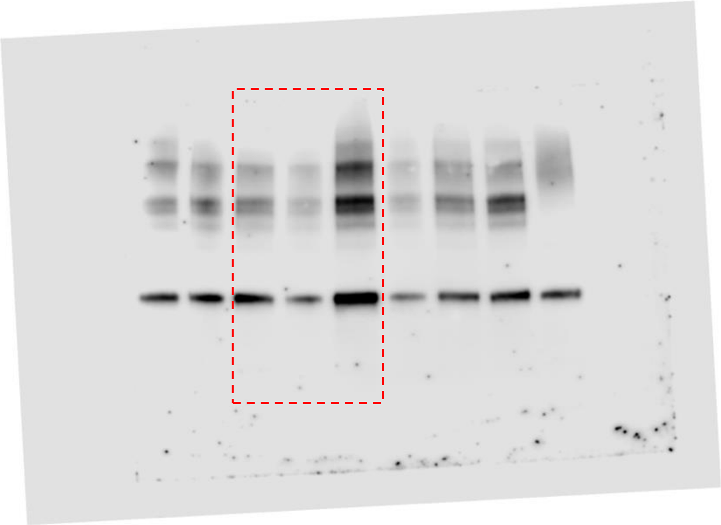

1G

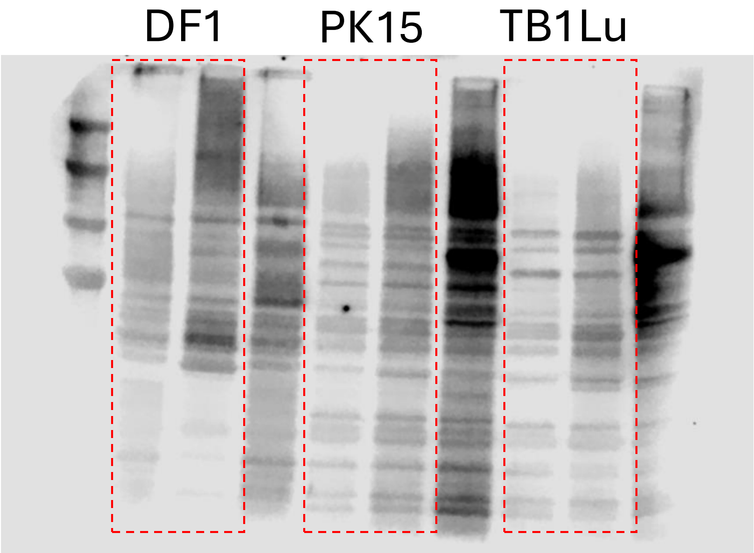


1H

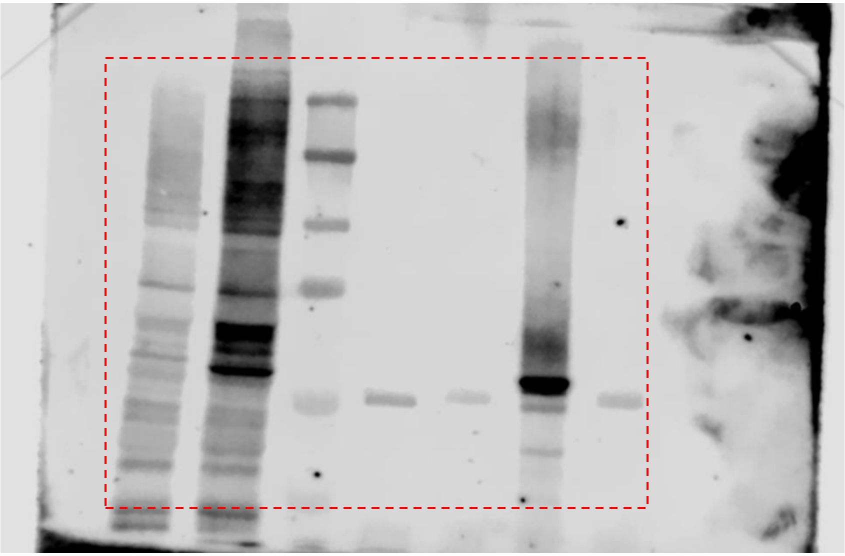

1I

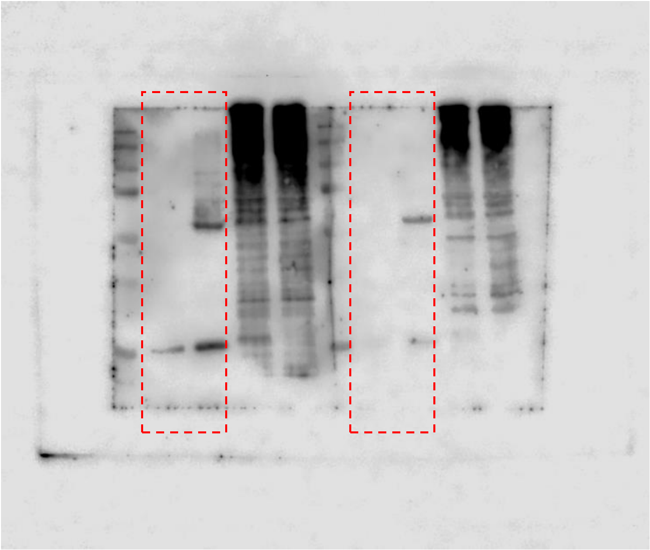

Figure 2F

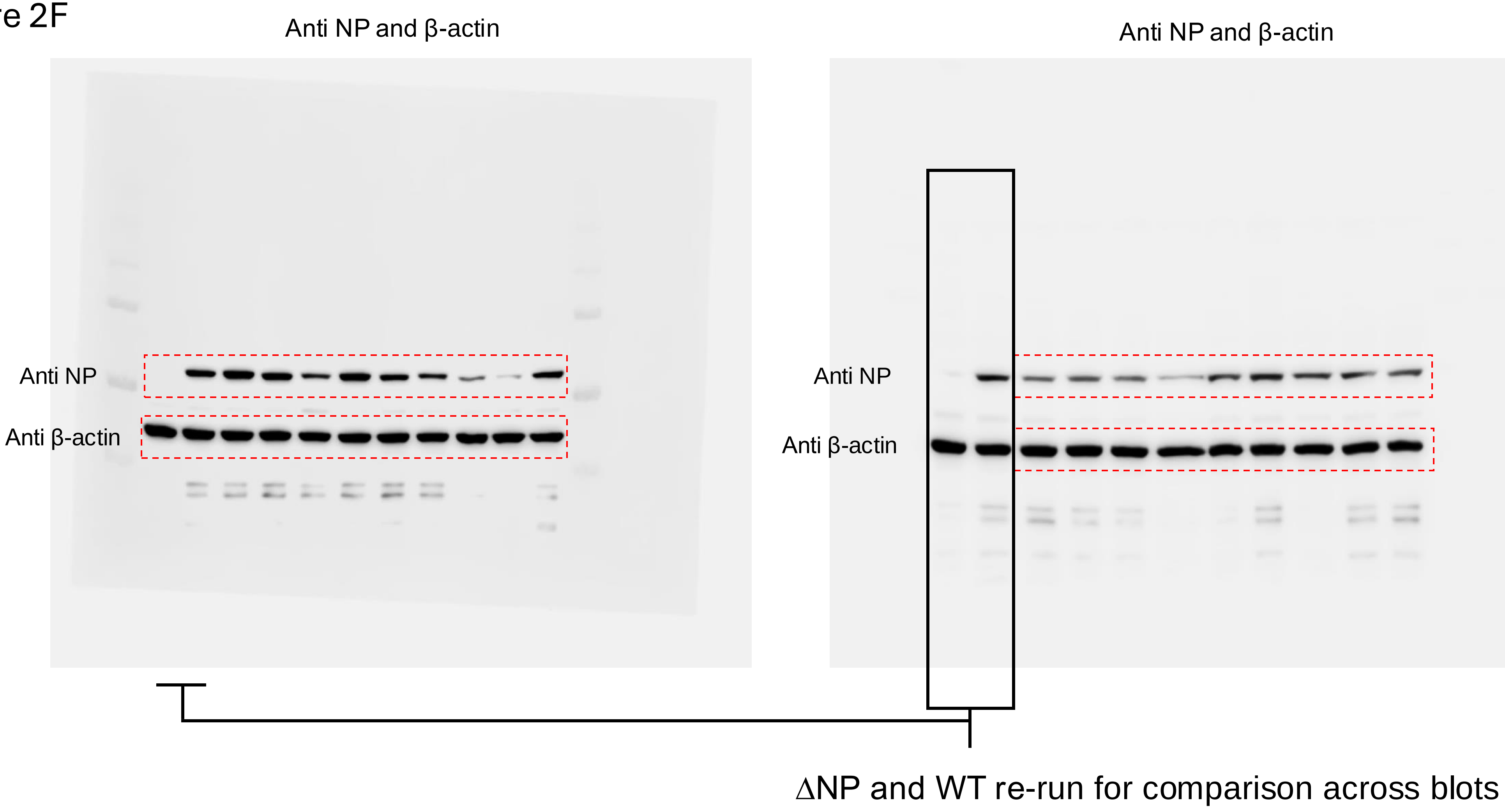

Anti NP and β-actin
Anti NP and β-actin
Anti NP
Anti β-actin
Anti NP
Anti β-actin
ΔNP and WT re-run for comparison across blots

### Figure 3A

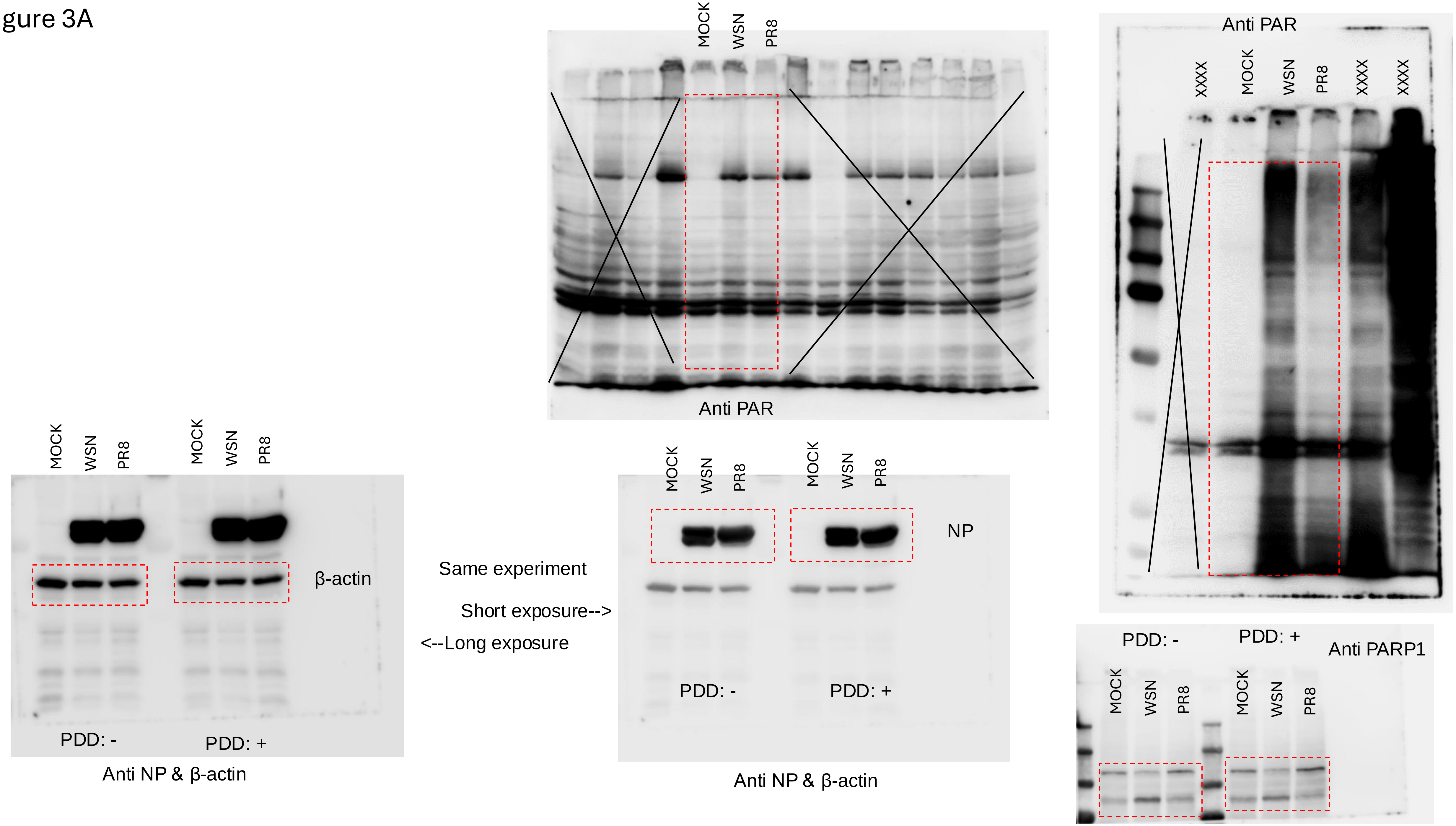

MOCK
WSN
PR8
Anti PAR
Anti PAR
XXXX
MOCK
WSN
PR8
XXXX
XXXX
MOCK
WSN
PR8
MOCK
WSN
PR8
β-actin
PDD: -
PDD: +
Anti NP & β-actin
Same experiment
Short exposure-->
<--Long exposure
MOCK
WSN
PR8
MOCK
WSN
PR8
NP
PDD: -
PDD: +
Anti NP & β-actin
PDD: -
PDD: +
Anti PARP1
MOCK
WSN
PR8
MOCK
WSN
PR8

Figure 3B

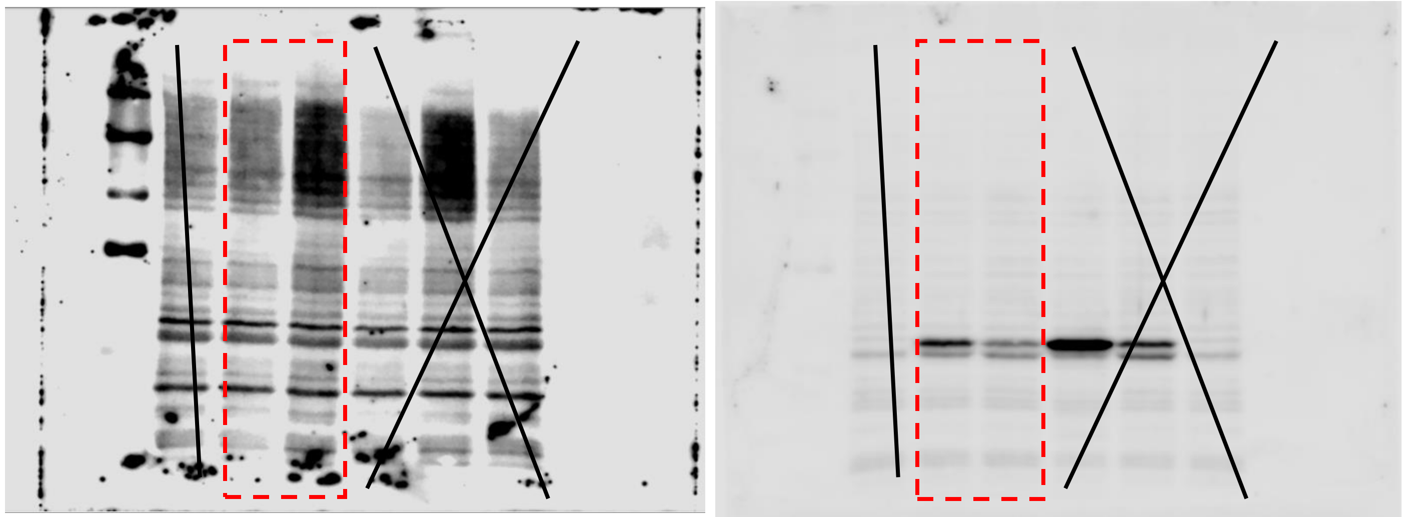

Figure 3C

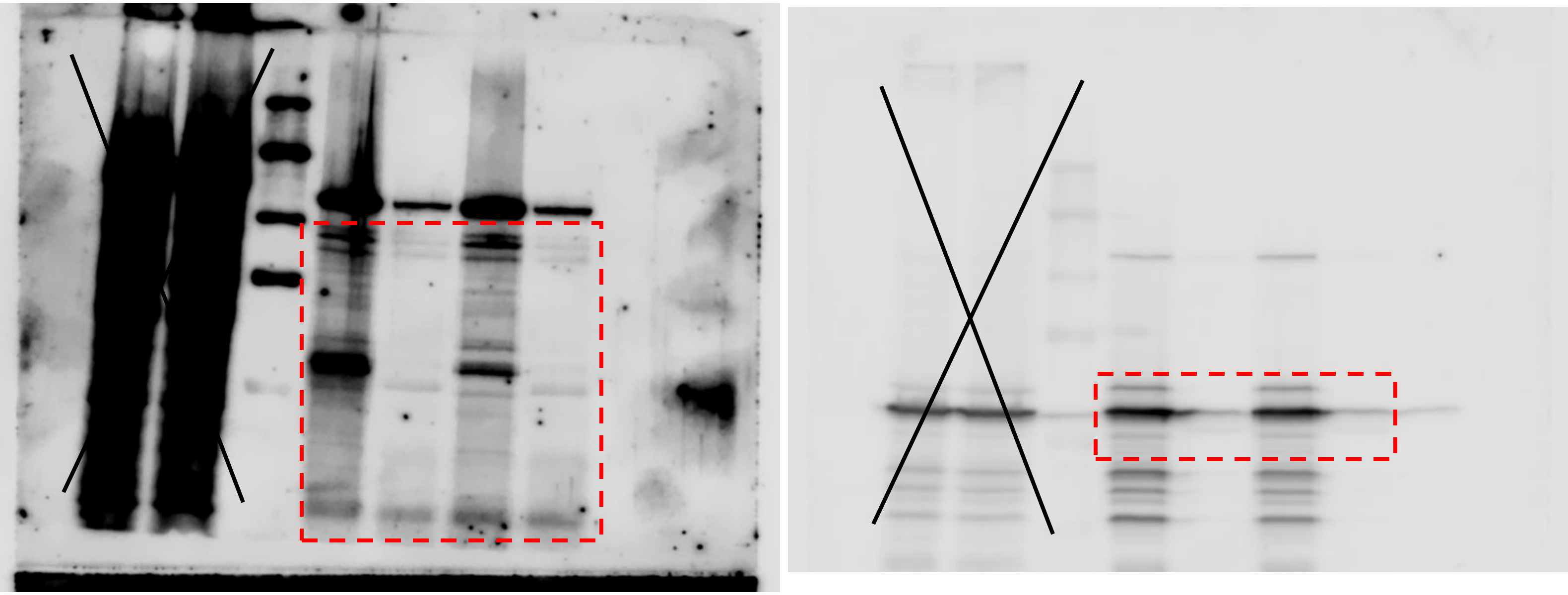

Figure 3D

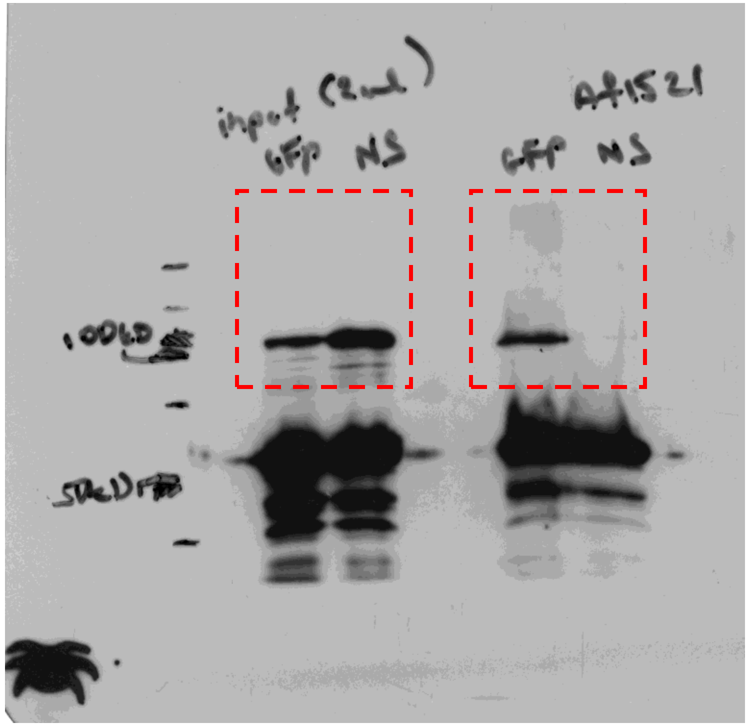

Figure 4A

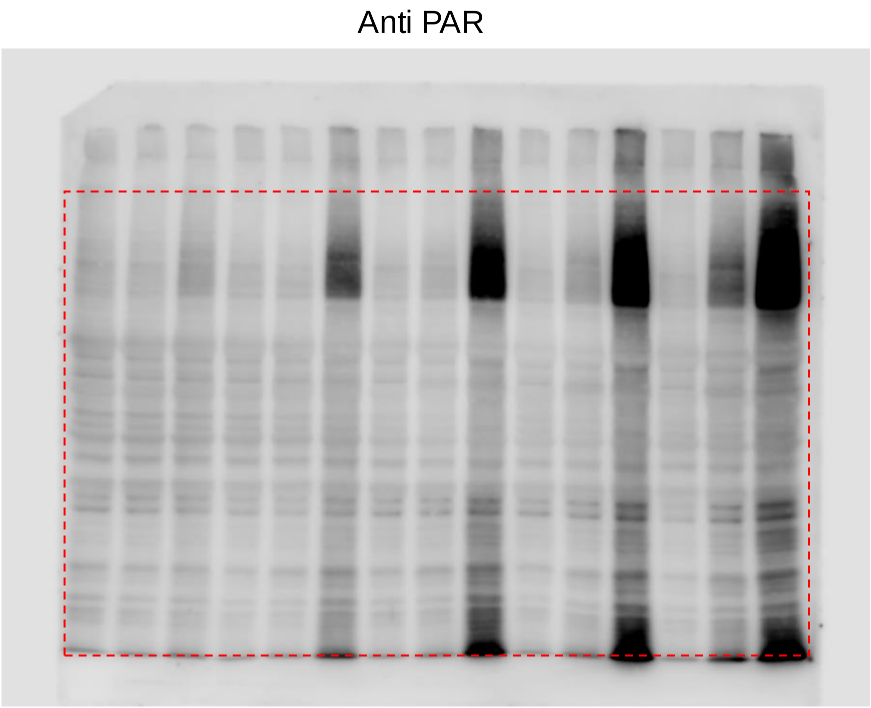


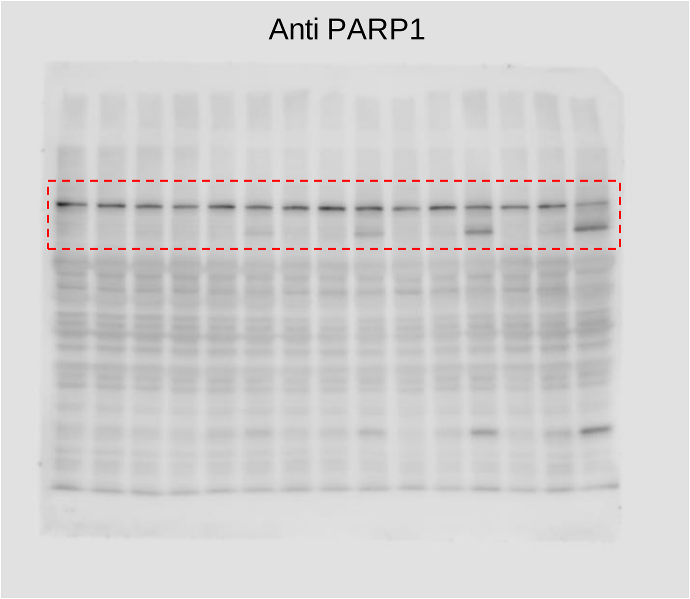


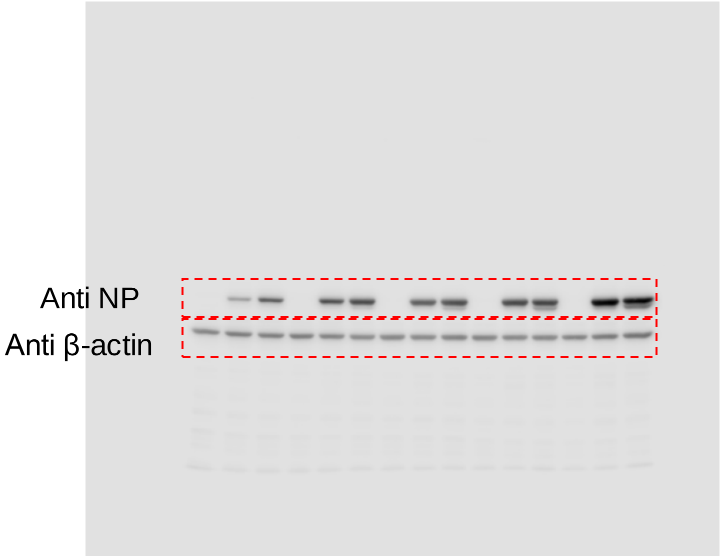

Figure 4B

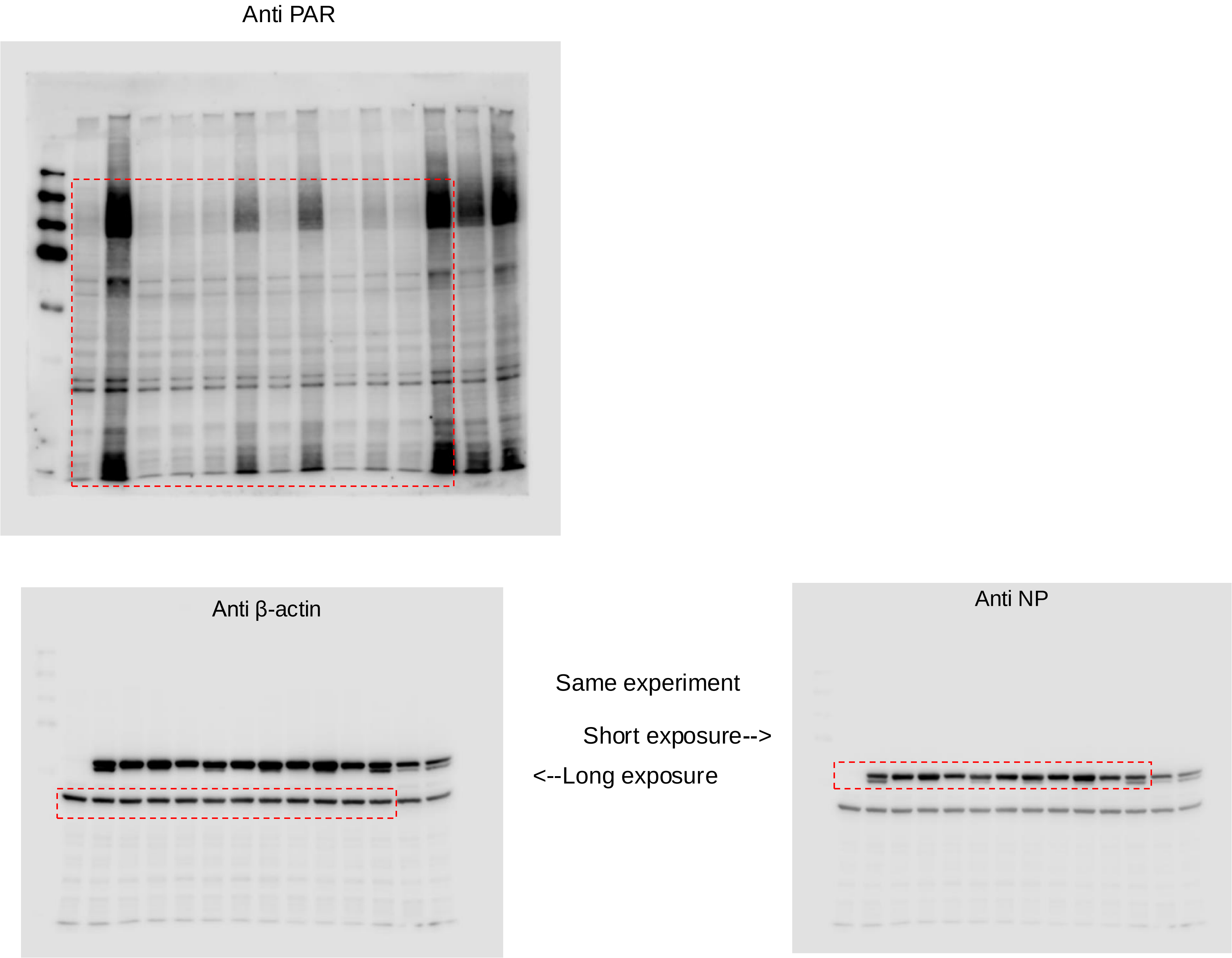
Anti PAR
Anti β-actin
Same experiment
Short exposure-->
<--Long exposure
Anti NP

### Figure 6A

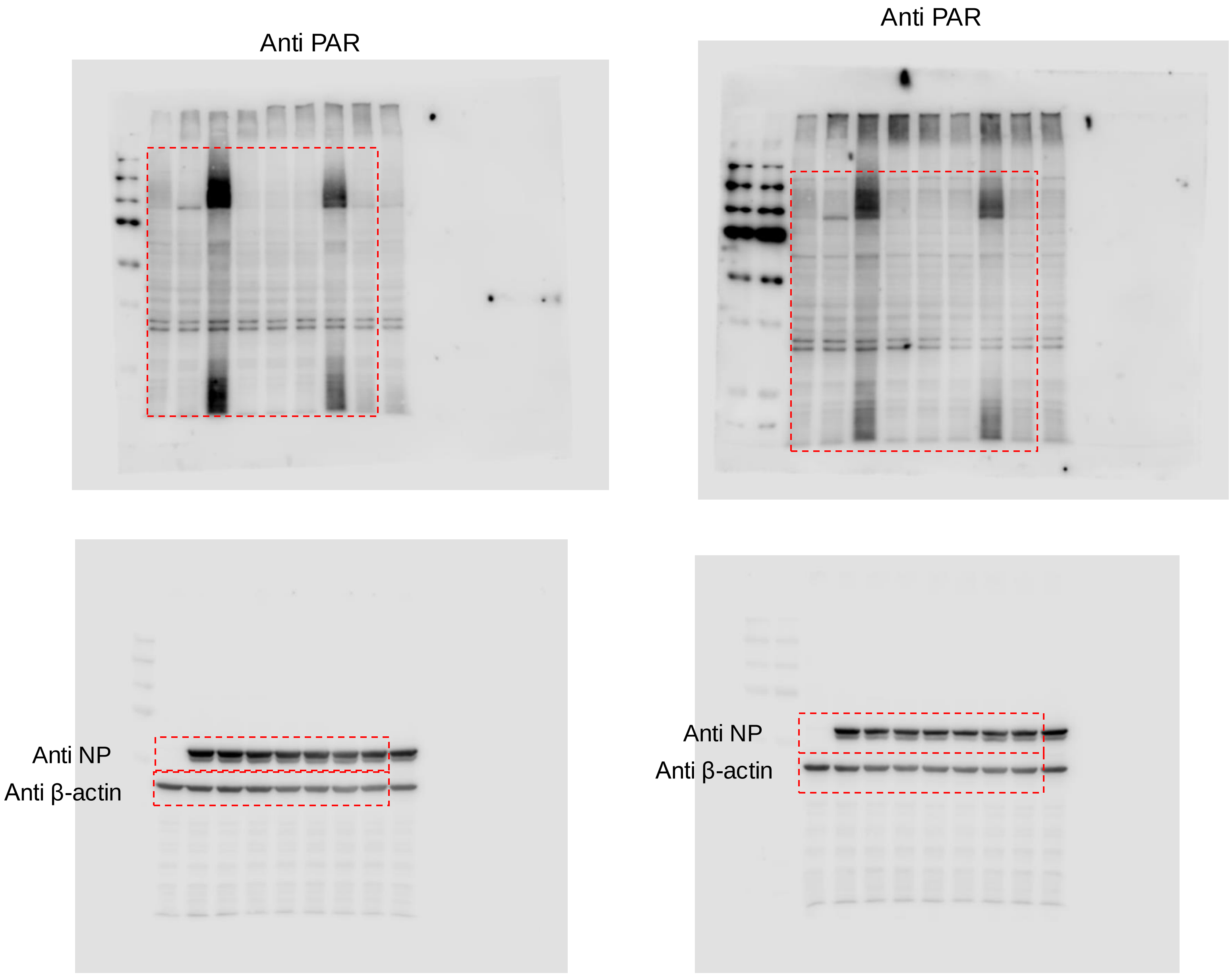

#### Figure 6B

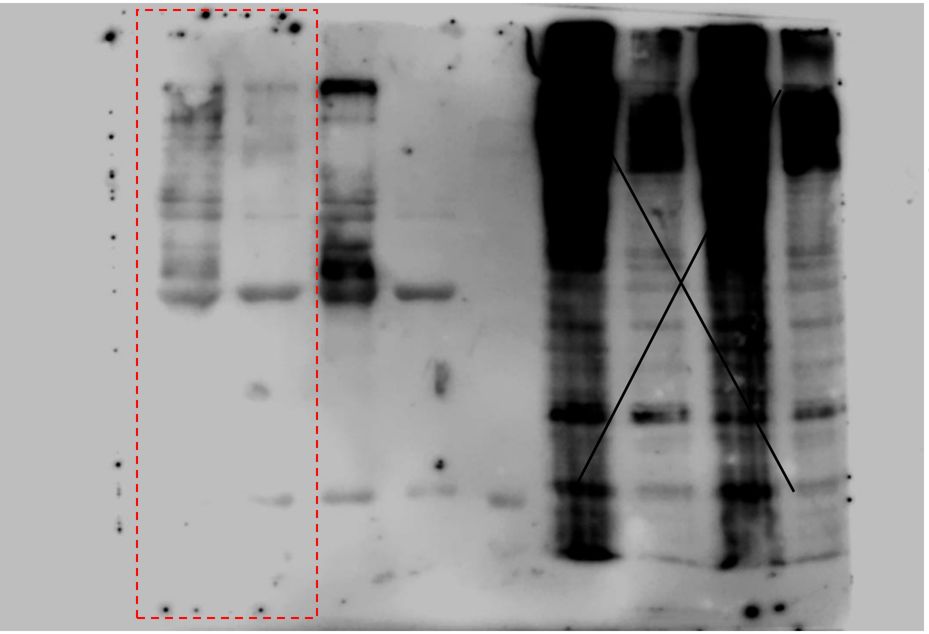

Same experiment

Short exposure-->

<--Long exposure

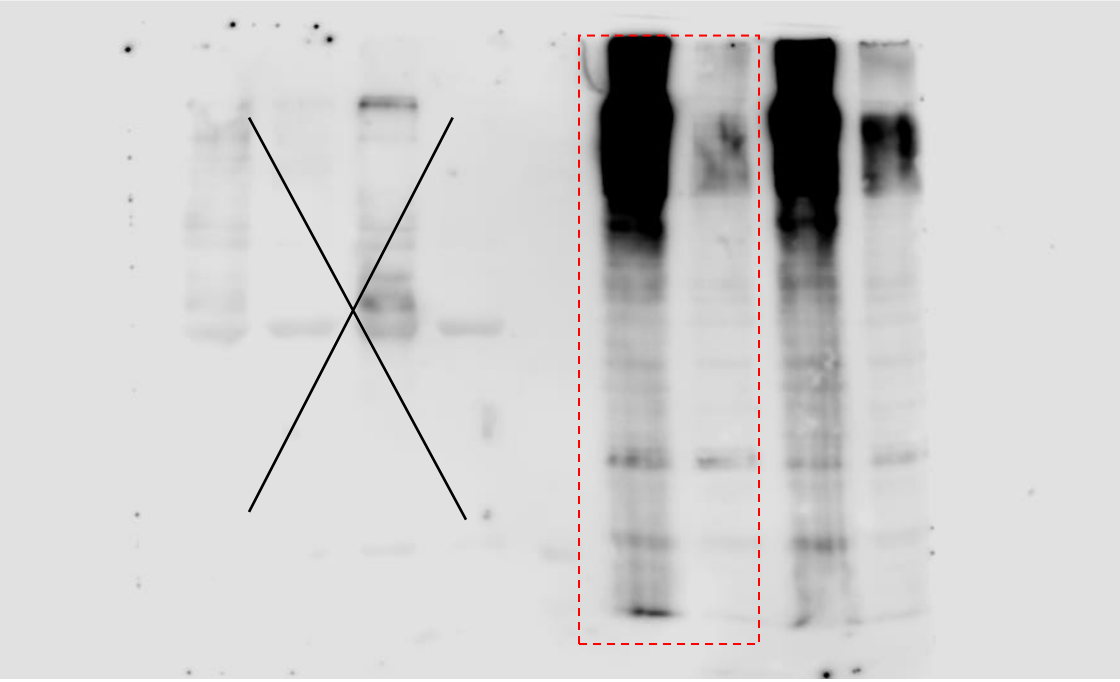

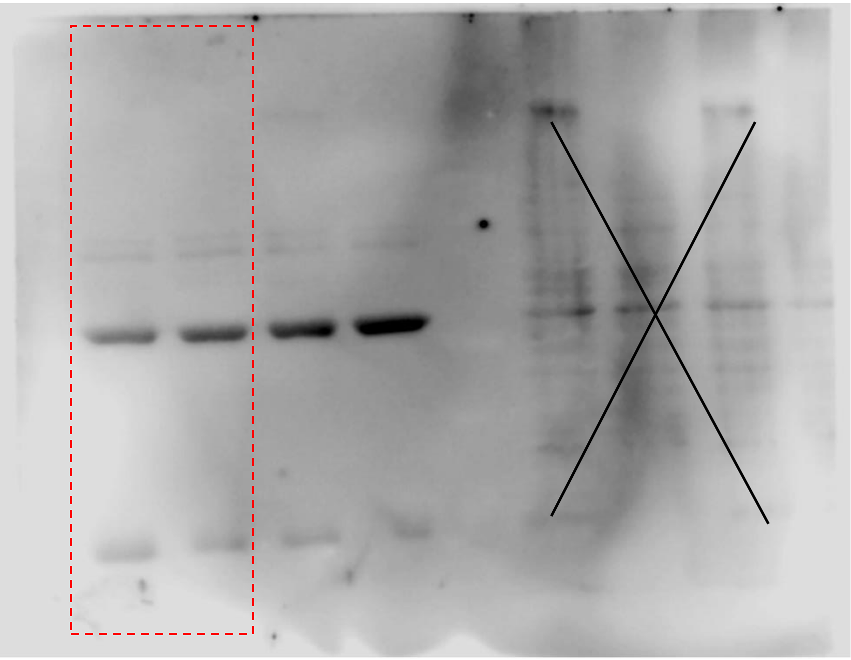

Figure 6C

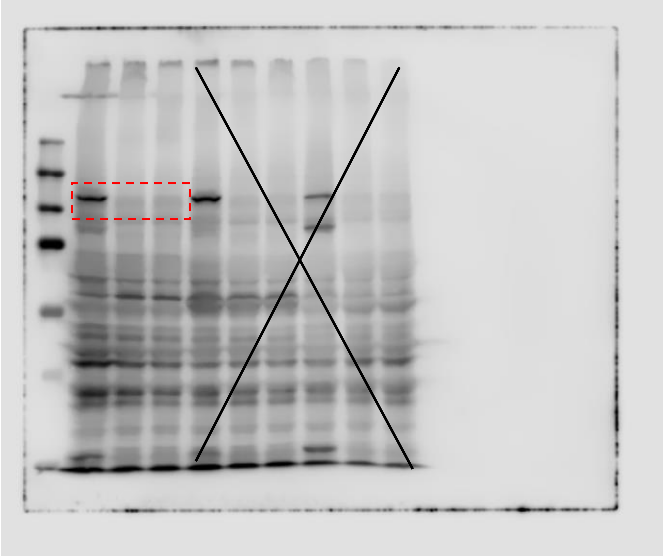


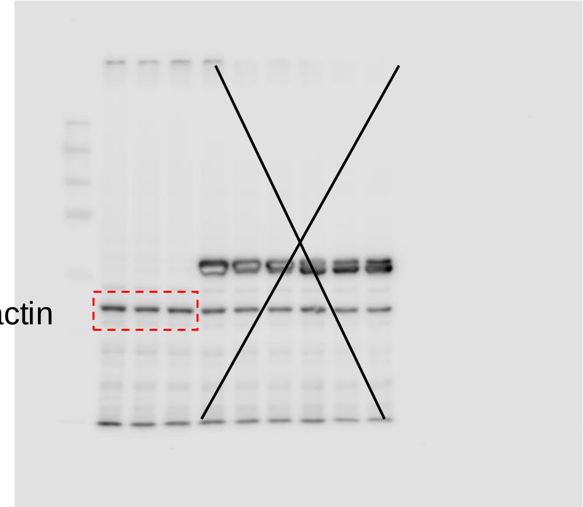

Figure 6D

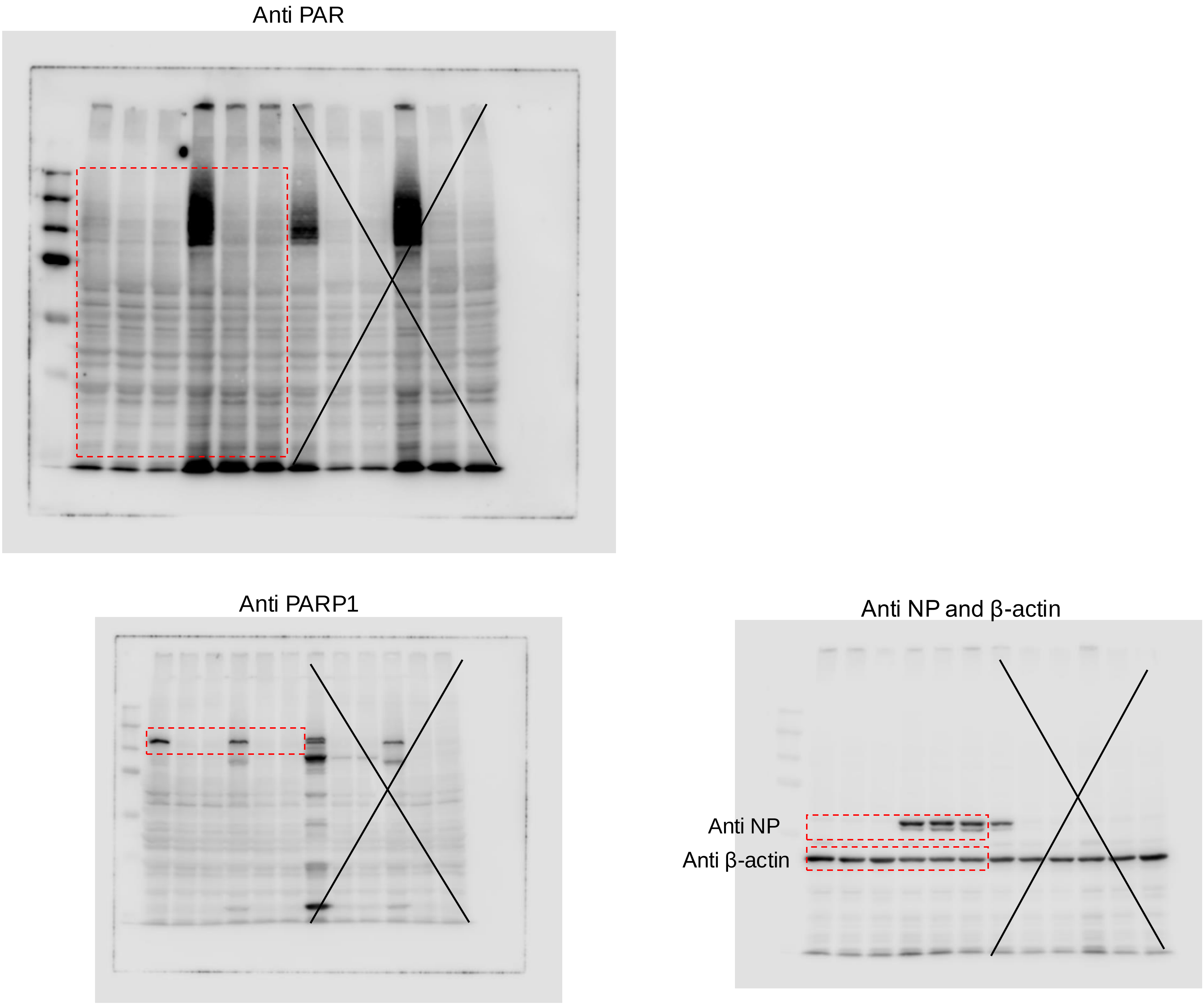

### Figure 6E

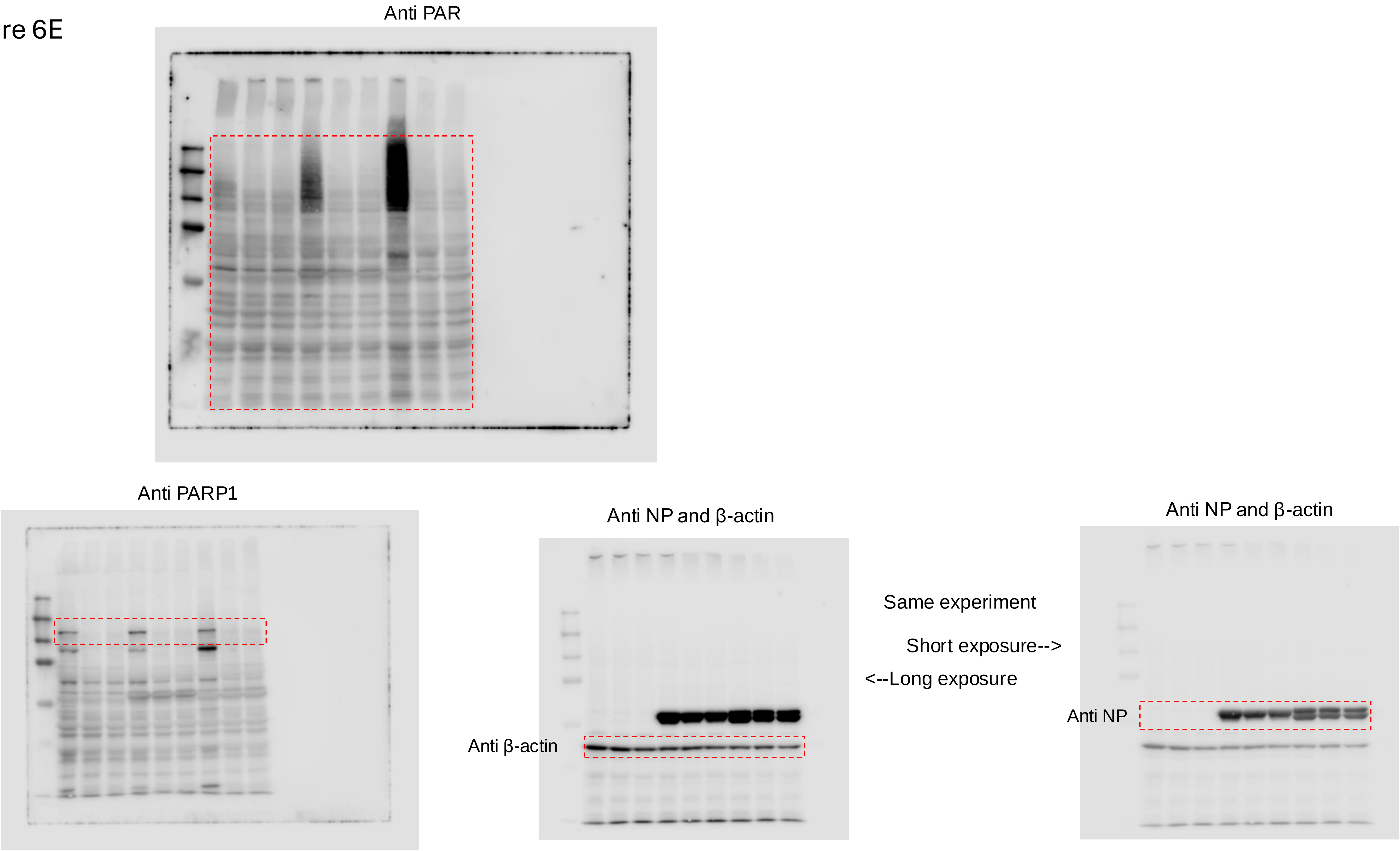
Anti PAR
Anti PARP1
Anti NP and β-actin
Anti β-actin
Same experiment
Short exposure-->
<--Long exposure
Anti NP and β-actin
Anti NP

### Figure 7A

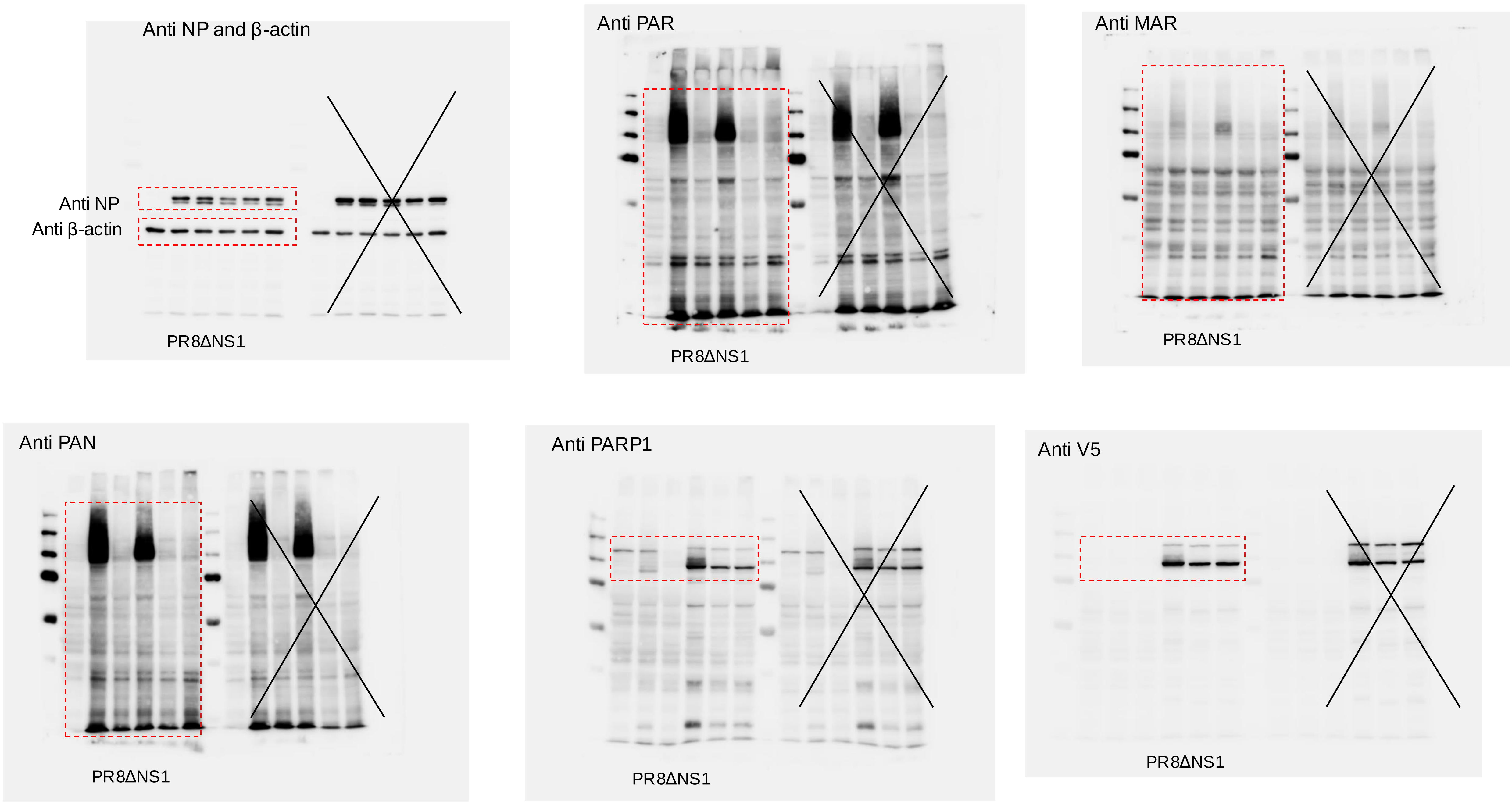
Anti NP and β-actin
Anti NP
Anti β-actin
PR8ΔNS1
Anti PAR
PR8ΔNS1
Anti MAR
PR8ΔNS1
Anti PAN
PR8ΔNS1
Anti PARP1
PR8ΔNS1
Anti V5
PR8ΔNS1
